# Antibody-mediated cellular responses are dysregulated in Multisystem Inflammatory Syndrome in Children (MIS-C)

**DOI:** 10.1101/2024.04.16.589585

**Authors:** Jenna K Dick, Jules A Sangala, Venkatramana D Krishna, Aaron Khaimraj, Lydia Hamel, Spencer M Erickson, Dustin Hicks, Yvette Soigner, Laura E Covill, Alexander Johnson, Michael J Ehrhardt, Keenan Ernst, Petter Brodin, Richard A Koup, Alka Khaitan, Carly Baehr, Beth K Thielen, Christine M Henzler, Caleb Skipper, Jeffrey S Miller, Yenan T Bryceson, Jianming Wu, Chandy C John, Angela Panoskaltsis-Mortari, Alberto Orioles, Marie E Steiner, Maxim C-J Cheeran, Marco Pravetoni, Geoffrey T Hart

## Abstract

Multisystem Inflammatory Syndrome in Children (MIS-C) is a severe complication of SARS-CoV-2 infection characterized by multi-organ involvement and inflammation. Testing of cellular function *ex vivo* to understand the aberrant immune response in MIS-C is limited. Despite strong antibody production in MIS-C, SARS-CoV-2 nucleic acid testing can remain positive for 4-6 weeks after infection. Therefore, we hypothesized that dysfunctional cell-mediated antibody responses downstream of antibody production may be responsible for delayed clearance of viral products in MIS-C. In MIS-C, monocytes were hyperfunctional for phagocytosis and cytokine production, while natural killer (NK) cells were hypofunctional for both killing and cytokine production. The decreased NK cell cytotoxicity correlated with an NK exhaustion marker signature and systemic IL-6 levels. Potentially providing a therapeutic option, cellular engagers of CD16 and SARS-CoV-2 proteins were found to rescue NK cell function *in vitro*. Together, our results reveal dysregulation in antibody-mediated cellular responses unique to MIS-C that likely contribute to the immune pathology of this disease.

**Summary:** MIS-C is a severe complication of SARS-CoV-2 infection characterized by multi-organ involvement and inflammation. Limited studies tested cellular function *ex vivo* to understand the aberrant immune response in MIS-C. We found dysregulation in antibody-mediated cellular responses unique to MIS-C that likely contribute to the immune pathology of this disease

Graphical abstract

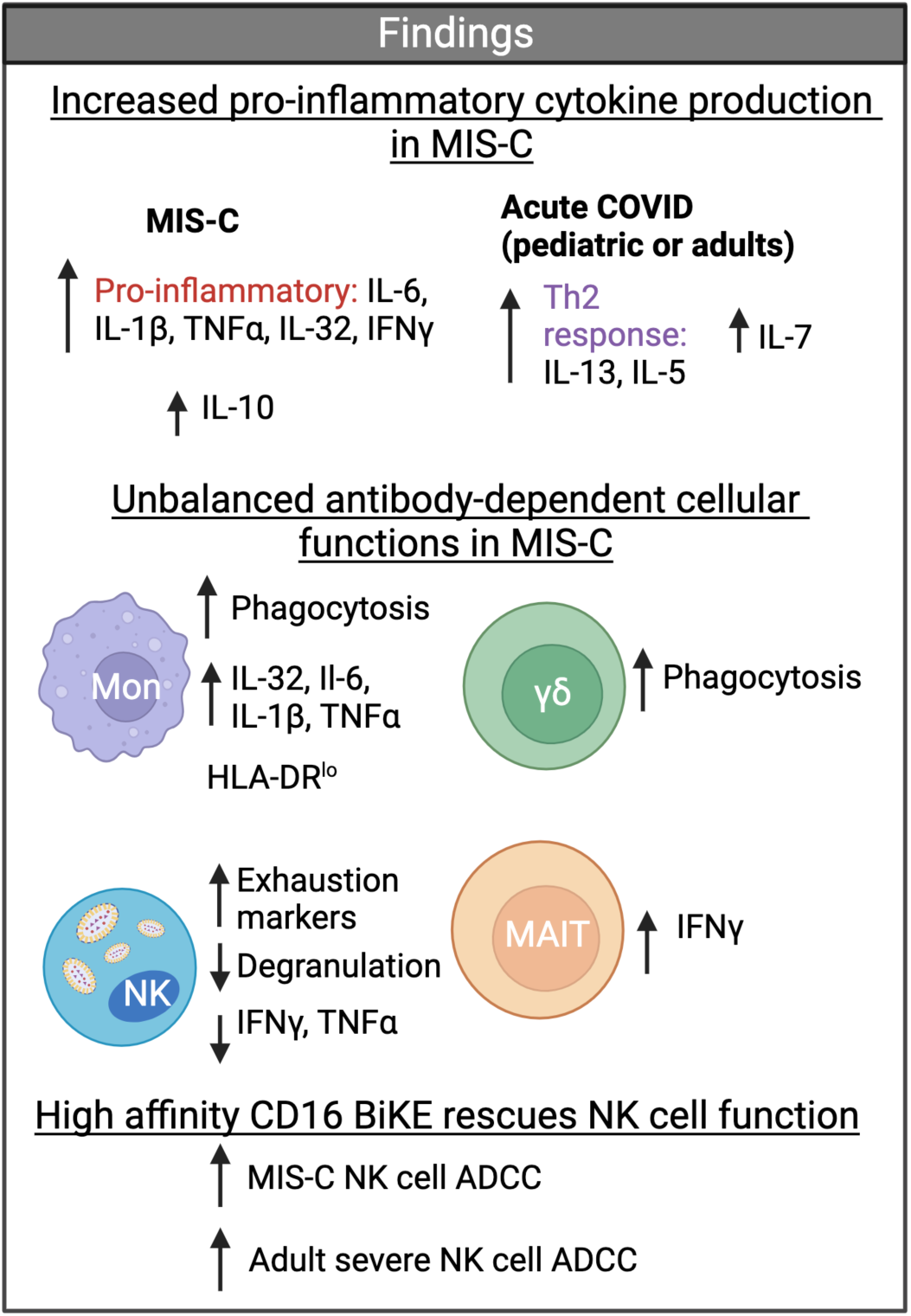

## Introduction

Children experience much lower rates of severe SARS-CoV-2 disease than adults. However, some children develop a rare and severe form of the disease known as Multisystem Inflammatory Syndrome in Children (MIS-C). Over half of children with MIS-C present to the intensive care unit (ICU) 4-6 weeks after initial SARS-CoV-2 infection (Cui et al., 2021; Patel, 2022; Vogel et al., 2021). Hallmark symptoms of MIS-C include a prolonged high fever, rash, gastrointestinal symptoms, cardiovascular dysfunction, and signs of inflammation (Feldstein et al., 2020; Hoste et al., 2021; Sönmez et al., 2022; Whittaker et al., 2020). Early recognition and treatment of MIS-C are essential to prevent serious complication and death. Children with MIS-C typically exhibit clinical improvement when treated with intravenous immunoglobulin (IVIG), steroids, and in some cases, Anakinra, an IL-1R antagonist; these treatments suggest a systemic immune dysregulation for disease pathogenesis (Akkoyun et al., 2023).

Understanding the immune processes associated with the development and pathogenesis of MIS-C is essential to develop better prevention and treatment strategies. Initially, MIS-C was thought to be an atypical form of Kawasaki Disease (KD), which is a hyperinflammatory syndrome associated with vasculitis and heart disease in children under the age of 5 years, hypothesized to be triggered by an unknown virus. However, there is now epidemiological and virological evidence indicating that SARS-CoV-2 is the specific trigger for MIS-C, and MIS-C is distinct from KD (Noval Rivas and Arditi, 2023; Rowley, 2020; Sharma et al., 2021; Vogel et al., 2021; Yasuhara et al., 2021). The inflammatory profiles of MIS-C resemble macrophage activation syndrome (MAS), an inflammatory syndrome often triggered by a preceding infection. MAS is characterized by uncontrolled activation and proliferation of T cells and macrophages accompanied by hypofunctional cytotoxic cells (Grom et al., 2003; Sacco et al., 2022; Sharma et al., 2021; Yasuhara et al., 2021). Distinct from MAS, MIS-C induces acute gastrointestinal symptoms and more prominent myocardial dysfunction. Together, these data suggest MIS-C is similar but distinct from KD and MAS (Carter et al., 2020; Filippatos et al., 2023; Lazova et al., 2022; Lee et al., 2020; Sacco et al., 2022).

Due to the limited availability of clinical research samples, there is minimal understanding of the underlying cellular mechanisms responsible for MIS-C. Concerning anti-viral antibodies, children with MIS-C develop anti-SARS-CoV-2 antibodies that persist and neutralize effectively (Burbelo et al., 2022; Ramaswamy et al., 2021; Richardson et al., 2022; Rostad et al., 2022; Ullah et al., 2021b; Weisberg et al., 2021). Once virus-specific antibodies develop and bind to viral proteins, Fc receptor expressing cells can, in-turn, bind to the antibody Fc domain and clear the virus via antibody-mediated effector functions. Several studies have shown that Fc-mediated antibody effector functions play a role in determining the outcome of acute SARS-CoV-2 infection in both mice and humans (Adeniji et al., 2021; Bahnan et al., 2021; Chakraborty et al., 2022; Filippatos et al., 2023; Richardson et al., 2022; Schäfer et al., 2021; Ullah et al., 2021a; Ullah et al., 2021b; Winkler et al., 2021; Yamin et al., 2021; Zhang et al., 2023; Zhang et al., 2022). However, it is unknown *how* innate immune cells that express Fc receptors respond to opsonized virus or viral producing cells in MIS-C.

Many immune cells that interact with antibodies via Fc receptors to clear viral particles have strong antibody-mediated effector functions, so they have the potential to be beneficial or pathogenic in the response to SARS-CoV-2. For example, monocytes can perform antibody-dependent cellular phagocytosis (ADCP) to clear virus or virally infected cells; too low of ADCP activity could contribute to outgrowth of SARS-CoV-2 viruses (Tay et al., 2019), whereas too strong of ADCP cytokine production could contribute to pathogenic inflammation (Andersson, 2021; Crayne et al., 2019). Gene signatures of monocytes in adults with severe acute COVID-19 infection and in children with MIS-C suggest myeloid activation (Idiz et al., 2022; Junqueira et al., 2022; Knoll et al., 2021). However, no *ex vivo* cellular functional studies have been performed on the innate cell compartment in MIS-C to see determine if ADCP functions correlate with pathology.

Additionally, NK cells mediate antibody-dependent cellular cytotoxicity (ADCC) through FcγRIIIA (CD16). Like monocytes, the strength of the NK cell response could result in increased viral load or pathogenic inflammation if the ADCC is too weak or strong, respectively. Of note, NK cells can kill cells without the need for antibody opsonization through a process called natural cytotoxicity (NC). Through this mechanism NK cells can target virally infected cells and can also kill other stressed cells. However, this mechanism could become dysfunctional if NK cells are exhausted. NK cell exhaustion leads to NK cell dysfunction and is identified by changes in marker expression, such as PD-1, TIGIT, CTLA-4, and TIM-3. Increased proportions of some of these markers have been associated with decreased function, and others with increased NK cell function depending on the context (Bi and Tian, 2017; Jia et al., 2023; Moebius et al., 2020; Rakova et al., 2021). These complex phenotypes, along with mechanisms on how NK cells become exhausted and if it can be reversed, are still being determined. NK cell function has not been studied in MIS-C.

Given that children with MIS-C have high levels of neutralizing antibodies but remain SARS-CoV-2 PCR positive for weeks after initial infection, we hypothesized that antibody-mediated cellular functions in MIS-C were dysregulated, leading to loss of homeostasis, prolonged infection, and subsequent pathology. To investigate the unique cellular antibody-mediated functions in children with MIS-C, we collected samples from children with MIS-C along with pediatric and adult individuals with a clinical spectrum of COVID-19 symptoms and uninfected controls. This enabled the comparison of Fc-mediated antibody functions across ages and clinical symptoms. We found that children with MIS-C had a neutralizing antibody response and a highly inflammatory cytokine milieu, consistent with previous studies (Consiglio et al., 2020; Gurlevik et al., 2022; Ravichandran et al., 2021; Rybkina et al., 2023; Thiriard et al., 2023). Leveraging the PBMC samples collected, we performed flow cytometric immunophenotyping and *ex vivo* functional assays with leukocytes expressing Fc receptors. We found dysregulated antibody-mediated cellular responses. Specifically, monocytes had enhanced ADCP and cytokine production in MIS-C compared to control groups. In contrast, NK cells produced fewer cytokines, degranulated less in an ADCC assay, and had significantly lower levels of perforin relative to control groups. We show that an NK exhaustion marker signature and more systemic IL-6 correlated with decreased NK cell function. Using novel bi and tri-specific reagents that target CD16 on NK cells with high affinity, we rescued the hypofunctional antibody-mediated NK cell function. Our results highlight that the cellular response to antibodies can be dysregulated during MIS-C and that regulation across many cell types is likely necessary to both decrease viral load and limit pathology.

## Results

### Clinical groups of MIS-C and COVID-19

To study the innate responses of children with MIS-C relative to other severities of acute SARS-CoV-2 infection, we analyzed both pediatric and adult subjects. There were a total of 66 infected subjects and 21 uninfected subjects categorized into MIS-C or one of 5 control groups: [1] children diagnosed with MIS-C based on the CDC definition: <21 years old, fever of >38 °C for >24 hrs, laboratory evidence of inflammation, hospital admission, multi-organ involvement, no alternative plausible diagnosis, and positive SARS-CoV-2 serology (Farooqi et al., 2021; Health, 2024) (n=14); [2] children <21 years old infected with SARS-CoV-2 acutely who did not develop MIS-C and were characterized as having severe infection (n=16); [3] children <21 years old infected with SARS-CoV-2 acutely who did not develop MIS-C and were characterized as having moderate infection (n=7); [4] children <21 years old that were infected with SARS-CoV-2 but were asymptomatic (n=15); [5] seronegative children <21 years old who had no evidence of ongoing or previous SARS-CoV-2 infection or vaccination (n=21); [6] adults who had acute severe SARS-CoV-2 infection (n=14). Demographics of each group and a schematic of enrollment timing over the pandemic in relation to the predominant circulating SARS-CoV-2 variant are in Table 1 and Figure 1 respectively.

**Figure 1.**
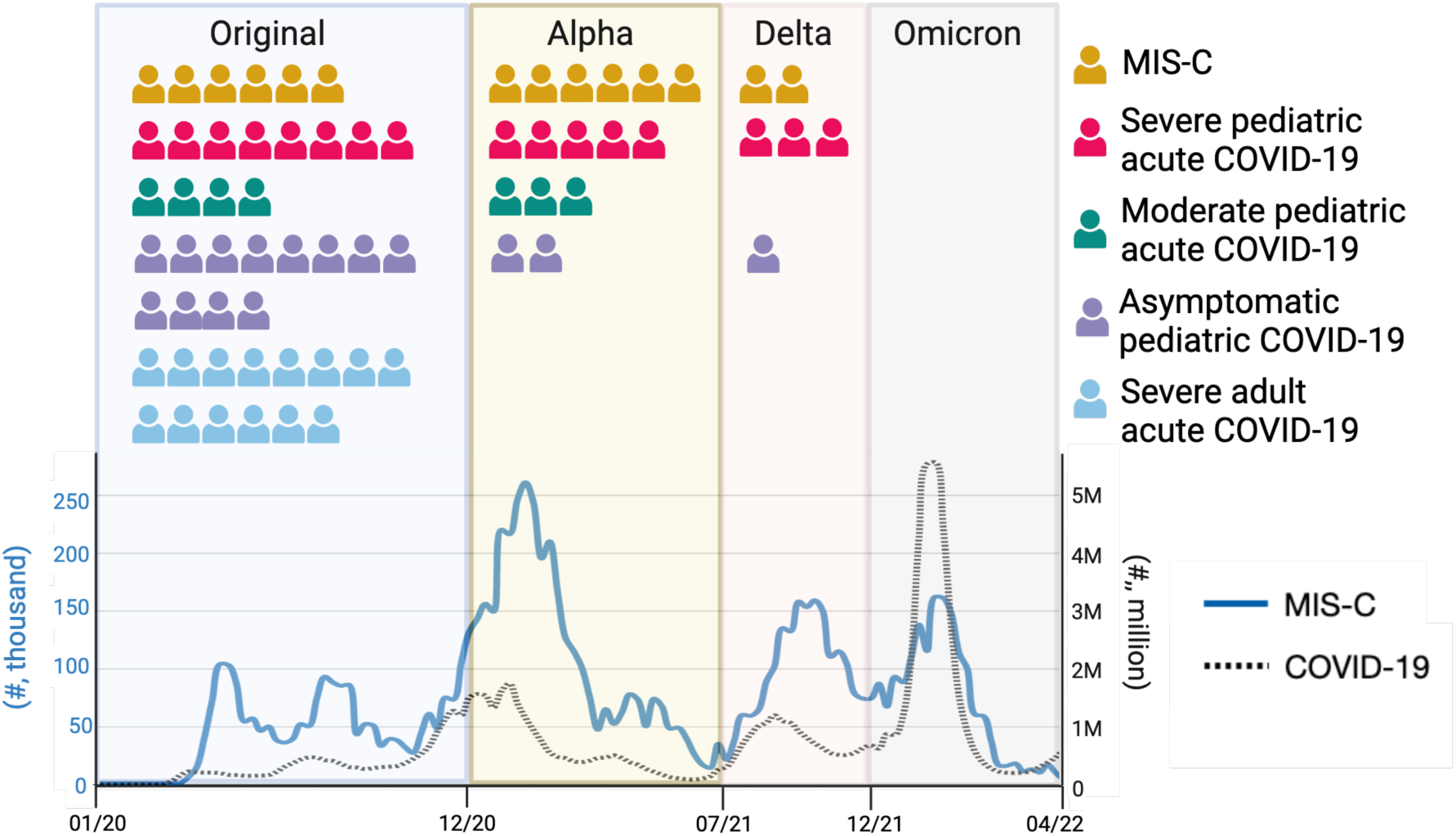
Study schematic of enrollment of each subject between 3/2020-12/2021 during the SARS-CoV-2 pandemic. Visual representation of each COVID-19+ subject enrolled and the predominant circulating variant when they were infected. The timing of acute COVID-19 infection in MIS-C patients is unknown and was presumed to be one month prior to MIS-C presentation (Feldstein et al., 2020). The weekly number of MIS-C cases is depicted in blue (left y axis) and the number of weekly COVID-19 cases overall is depicted in a black dotted line (right y axis). The case data was sourced from the Center for Disease Control. Figure based on Rybinka et al. (Rybkina et al., 2023).

**Table 1.**
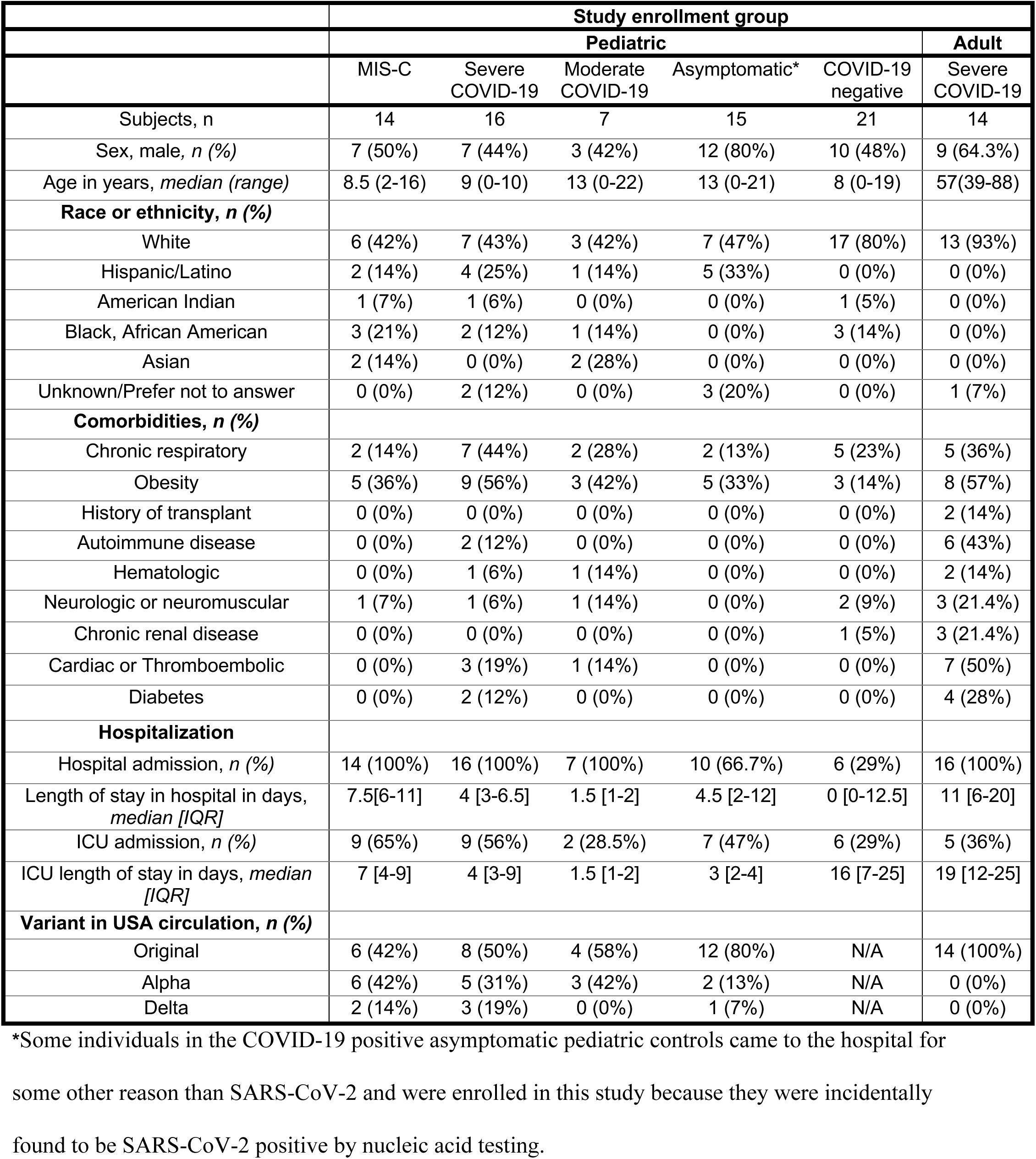
Demographics of patient groups.

|  | Study enrollment group |  |  |  |  |  |
| --- | --- | --- | --- | --- | --- | --- |
|  | Pediatric |  |  |  |  | Adult |
|  | MIS-C | Severe COVID-19 | Moderate COVID-19 | Asymptomatic* | COVID-19 negative | Severe COVID-19 |
| Subjects, n | 14 | 16 | 7 | 15 | 21 | 14 |
| Sex, male, *n* (%) | 7 (50%) | 7 (44%) | 3 (42%) | 12 (80%) | 10 (48%) | 9 (64.3%) |
| Age in years, *median (range)* | 8.5 (2-16) | 9 (0-10) | 13 (0-22) | 13 (0-21) | 8 (0-19) | 57(39-88) |
| **Race or ethnicity, *n* (%)** |  |  |  |  |  |  |
| White | 6 (42%) | 7 (43%) | 3 (42%) | 7 (47%) | 17 (80%) | 13 (93%) |
| Hispanic/Latino | 2 (14%) | 4 (25%) | 1 (14%) | 5 (33%) | 0 (0%) | 0 (0%) |
| American Indian | 1 (7%) | 1 (6%) | 0 (0%) | 0 (0%) | 1 (5%) | 0 (0%) |
| Black, African American | 3 (21%) | 2 (12%) | 1 (14%) | 0 (0%) | 3 (14%) | 0 (0%) |
| Asian | 2 (14%) | 0 (0%) | 2 (28%) | 0 (0%) | 0 (0%) | 0 (0%) |
| Unknown/Prefer not to answer | 0 (0%) | 2 (12%) | 0 (0%) | 3 (20%) | 0 (0%) | 1 (7%) |
| **Comorbidities, *n* (%)** |  |  |  |  |  |  |
| Chronic respiratory | 2 (14%) | 7 (44%) | 2 (28%) | 2 (13%) | 5 (23%) | 5 (36%) |
| Obesity | 5 (36%) | 9 (56%) | 3 (42%) | 5 (33%) | 3 (14%) | 8 (57%) |
| History of transplant | 0 (0%) | 0 (0%) | 0 (0%) | 0 (0%) | 0 (0%) | 2 (14%) |
| Autoimmune disease | 0 (0%) | 2 (12%) | 0 (0%) | 0 (0%) | 0 (0%) | 6 (43%) |
| Hematologic | 0 (0%) | 1 (6%) | 1 (14%) | 0 (0%) | 0 (0%) | 2 (14%) |
| Neurologic or neuromuscular | 1 (7%) | 1 (6%) | 1 (14%) | 0 (0%) | 2 (9%) | 3 (21.4%) |
| Chronic renal disease | 0 (0%) | 0 (0%) | 0 (0%) | 0 (0%) | 1 (5%) | 3 (21.4%) |
| Cardiac or Thromboembolic | 0 (0%) | 3 (19%) | 1 (14%) | 0 (0%) | 0 (0%) | 7 (50%) |
| Diabetes | 0 (0%) | 2 (12%) | 0 (0%) | 0 (0%) | 0 (0%) | 4 (28%) |
| **Hospitalization** |  |  |  |  |  |  |
| Hospital admission, *n* (%) | 14 (100%) | 16 (100%) | 7 (100%) | 10 (66.7%) | 6 (29%) | 16 (100%) |
| Length of stay in hospital in days, *median [IQR]* | 7.5[6-11] | 4 [3-6.5] | 1.5 [1-2] | 4.5 [2-12] | 0 [0-12.5] | 11 [6-20] |
| ICU admission, *n* (%) | 9 (65%) | 9 (56%) | 2 (28.5%) | 7 (47%) | 6 (29%) | 5 (36%) |
| ICU length of stay in days, *median [IQR]* | 7 [4-9] | 4 [3-9] | 1.5 [1-2] | 3 [2-4] | 16 [7-25] | 19 [12-25] |
| **Variant in USA circulation, *n* (%)** |  |  |  |  |  |  |
| Original | 6 (42%) | 8 (50%) | 4 (58%) | 12 (80%) | N/A | 14 (100%) |
| Alpha | 6 (42%) | 5 (31%) | 3 (42%) | 2 (13%) | N/A | 0 (0%) |
| Delta | 2 (14%) | 3 (19%) | 0 (0%) | 1 (7%) | N/A | 0 (0%) |

\*Some individuals in the COVID-19 positive asymptomatic pediatric controls came to the hospital for some other reason than SARS-CoV-2 and were enrolled in this study because they were incidentally found to be SARS-CoV-2 positive by nucleic acid testing.

In terms of demographic differences between MIS-C and control groups, the children who developed MIS-C, severe COVID-19 or moderate COVID-19 were more racially and ethnically diverse than any other groups. This is consistent with previous literature showing strong evidence of racial and ethnic disparities in SARS-CoV-2 severity (Hollis et al., 2021; Vahidy et al., 2020). In terms of clinical severity, the median hospital length of stay was 7.5 days for children with MIS-C, which was higher than any other pediatric group. Children with MIS-C were also more likely to be intubated than any other group (Table 2). In terms of treatment, most children with MIS-C received intravenous immunoglobulin (IVIG) (71%) while in the other groups, very few patients received IVIG. Of note, the IVIG was made before the COVID-19 pandemic started and did not contain SARS-CoV-2 specific antibodies. Also, no children with MIS-C received convalescent plasma (Table 2). With these groups, we sought to determine distinct immunological differences in children with MIS-C.

**Table 2.**
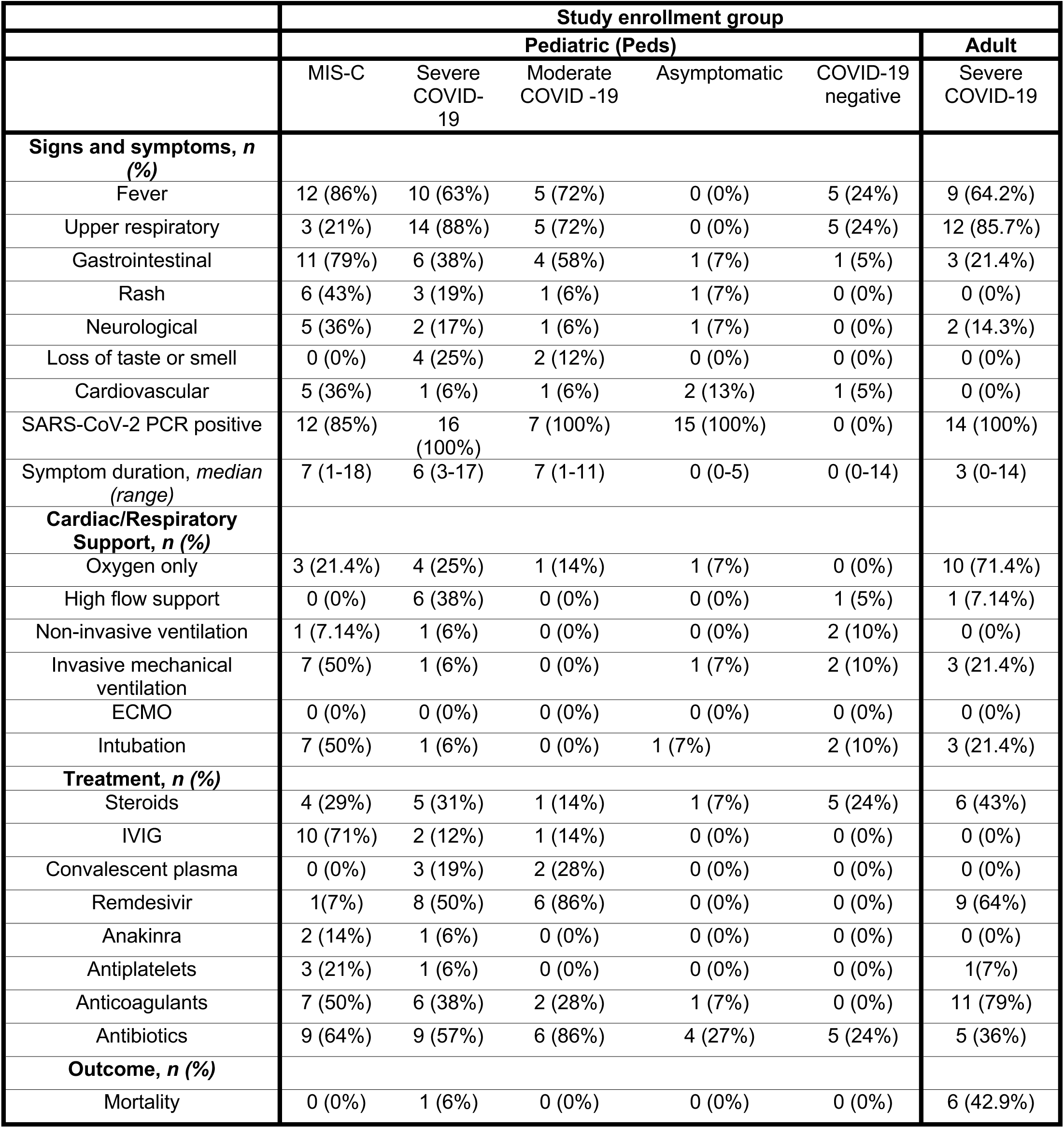
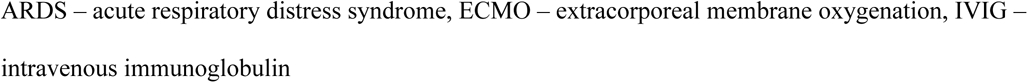
Clinical characteristics of patient groups.

|  | Study enrollment group |  |  |  |  |  |
| --- | --- | --- | --- | --- | --- | --- |
|  | Pediatric (Peds) |  |  |  |  | Adult |
|  | MIS-C | Severe COVID-19 | Moderate COVID -19 | Asymptomatic | COVID-19 negative | Severe COVID-19 |
| **Signs and symptoms, *n* (%)** |  |  |  |  |  |  |
| Fever | 12 (86%) | 10 (63%) | 5 (72%) | 0 (0%) | 5 (24%) | 9 (64.2%) |
| Upper respiratory | 3 (21%) | 14 (88%) | 5 (72%) | 0 (0%) | 5 (24%) | 12 (85.7%) |
| Gastrointestinal | 11 (79%) | 6 (38%) | 4 (58%) | 1 (7%) | 1 (5%) | 3 (21.4%) |
| Rash | 6 (43%) | 3 (19%) | 1 (6%) | 1 (7%) | 0 (0%) | 0 (0%) |
| Neurological | 5 (36%) | 2 (17%) | 1 (6%) | 1 (7%) | 0 (0%) | 2 (14.3%) |
| Loss of taste or smell | 0 (0%) | 4 (25%) | 2 (12%) | 0 (0%) | 0 (0%) | 0 (0%) |
| Cardiovascular | 5 (36%) | 1 (6%) | 1 (6%) | 2 (13%) | 1 (5%) | 0 (0%) |
| SARS-CoV-2 PCR positive | 12 (85%) | 16 (100%) | 7 (100%) | 15 (100%) | 0 (0%) | 14 (100%) |
| Symptom duration, *median (range)* | 7 (1-18) | 6 (3-17) | 7 (1-11) | 0 (0-5) | 0 (0-14) | 3 (0-14) |
| **Cardiac/Respiratory Support, *n* (%)** |  |  |  |  |  |  |
| Oxygen only | 3 (21.4%) | 4 (25%) | 1 (14%) | 1 (7%) | 0 (0%) | 10 (71.4%) |
| High flow support | 0 (0%) | 6 (38%) | 0 (0%) | 0 (0%) | 1 (5%) | 1 (7.14%) |
| Non-invasive ventilation | 1 (7.14%) | 1 (6%) | 0 (0%) | 0 (0%) | 2 (10%) | 0 (0%) |
| Invasive mechanical ventilation | 7 (50%) | 1 (6%) | 0 (0%) | 1 (7%) | 2 (10%) | 3 (21.4%) |
| ECMO | 0 (0%) | 0 (0%) | 0 (0%) | 0 (0%) | 0 (0%) | 0 (0%) |
| Intubation | 7 (50%) | 1 (6%) | 0 (0%) | 1 (7%) | 2 (10%) | 3 (21.4%) |
| **Treatment, *n* (%)** |  |  |  |  |  |  |
| Steroids | 4 (29%) | 5 (31%) | 1 (14%) | 1 (7%) | 5 (24%) | 6 (43%) |
| IVIg | 10 (71%) | 2 (12%) | 1 (14%) | 0 (0%) | 0 (0%) | 0 (0%) |
| Convalescent plasma | 0 (0%) | 3 (19%) | 2 (28%) | 0 (0%) | 0 (0%) | 0 (0%) |
| Remdesivir | 1(7%) | 8 (50%) | 6 (86%) | 0 (0%) | 0 (0%) | 9 (64%) |
| Anakinra | 2 (14%) | 1 (6%) | 0 (0%) | 0 (0%) | 0 (0%) | 0 (0%) |
| Antiplatelets | 3 (21%) | 1 (6%) | 0 (0%) | 0 (0%) | 0 (0%) | 1(7%) |
| Anticoagulants | 7 (50%) | 6 (38%) | 2 (28%) | 1 (7%) | 0 (0%) | 11 (79%) |
| Antibiotics | 9 (64%) | 9 (57%) | 6 (86%) | 4 (27%) | 5 (24%) | 5 (36%) |
| **Outcome, *n* (%)** |  |  |  |  |  |  |
| Mortality | 0 (0%) | 1 (6%) | 0 (0%) | 0 (0%) | 0 (0%) | 6 (42.9%) |

ARDS – acute respiratory distress syndrome, ECMO – extracorporeal membrane oxygenation, IVIG – intravenous immunoglobulin

### Children with MIS-C have strong antibody responses and increased pro-inflammatory cytokines compared to acute COVID-19 controls

To investigate the roles of antibody dependent cellular cytotoxicity (ADCC) and antibody dependent cellular phagocytosis (ADCP) in MIS-C, we initially assessed total antibody levels in each group. Most studies have measured Spike or nucleocapsid antibody levels. The Spike protein has two subunits, Spike 1 and Spike 2 (Fig. S1A) and Spike 1 is required for viral entry and is the target of neutralizing antibodies. Nucleocapsid protein generates one of the strongest antibody responses, its sequence is relatively less variable than Spike, and it is only present in natural infection and not in currently approved vaccines; this makes it a beneficial diagnostic tool (Matchett et al., 2021). We chose to enrich this data set by investigating antibodies and viral protein sequence variation against viral proteins available for antibody detection on the surface of the virus (Fig. S1A). Proteins expressed on the surface of SARS-CoV-2 are Spike, membrane, and envelope proteins. Antibodies to Spike 2 subunit protein and membrane (divided into M1 and M2) and envelope (E1) peptides expressed on the virus surface (Fig. S1A) are likely not involved in viral neutralization but may be involved in viral clearance through ADCC and ADCP. We found that children with MIS-C had a significant response (antibody levels were above baseline) to all available surface proteins including Spike 1 subunit (neutralizing) and non-neutralizing surface targets like the Spike 2 subunit protein and membrane (M1 and M2) and envelope (E1) peptides (Fig. S1 B-G). We also found that the variations in sequences of the non-neutralizing proteins and peptides (Spike 2, M1, M2, and E1) were less than that of Spike 1 over time (Fig. S2). We found an overall higher antibody response in the children with MIS-C, but this is consistent with those individuals having been infected multiple weeks longer than our controls.

A key hallmark of MIS-C and severe COVID-19 in adults is the elevation of pro-inflammatory cytokines (Consiglio et al., 2020; Diorio et al., 2021; Rybkina et al., 2023; Szabo et al., 2021). We therefore compared cytokine and chemokine profiles as measured by multiplex analysis at the group and individual subject level (Fig. S1I), and measured pan-IL-32 by ELISA. IL-32 is a proinflammatory cytokine only found in humans. It is primarily expressed by NK cells, monocytes, and T cells and is involved in inducing other pro-inflammatory cytokines, such as IL-8 and TNFα (Bergantini et al., 2022; Kim et al., 2005; Shim et al., 2022). We found that children with MIS-C exhibited distinct and significantly increased cytokine levels compared with other groups. These MIS-C-specific elevations includes higher levels of cytokines associated with cytokine storm such as IL-6, IL-1β, GM-CSF, IFNγ, and TNFα (Fig. 2B-F). Additionally, children with MIS-C had significantly higher levels of IL-32 compared to many other control groups (Fig. 2G). They also showed an increase in the immunosuppressive cytokine IL-10 (Fig. 1A and S1I) (Dhar et al., 2021; Newton et al., 2016). Those with acute COVID-19 infection showed higher levels of cytokines involved in Th2 responses, include IL-13 and IL-15. They also have an increase in IL-7. These results identify distinct cytokines and chemokines that are at higher levels in MIS-C as compared to other acute COVID-19 pediatric and adult responses.

**Figure 2.**
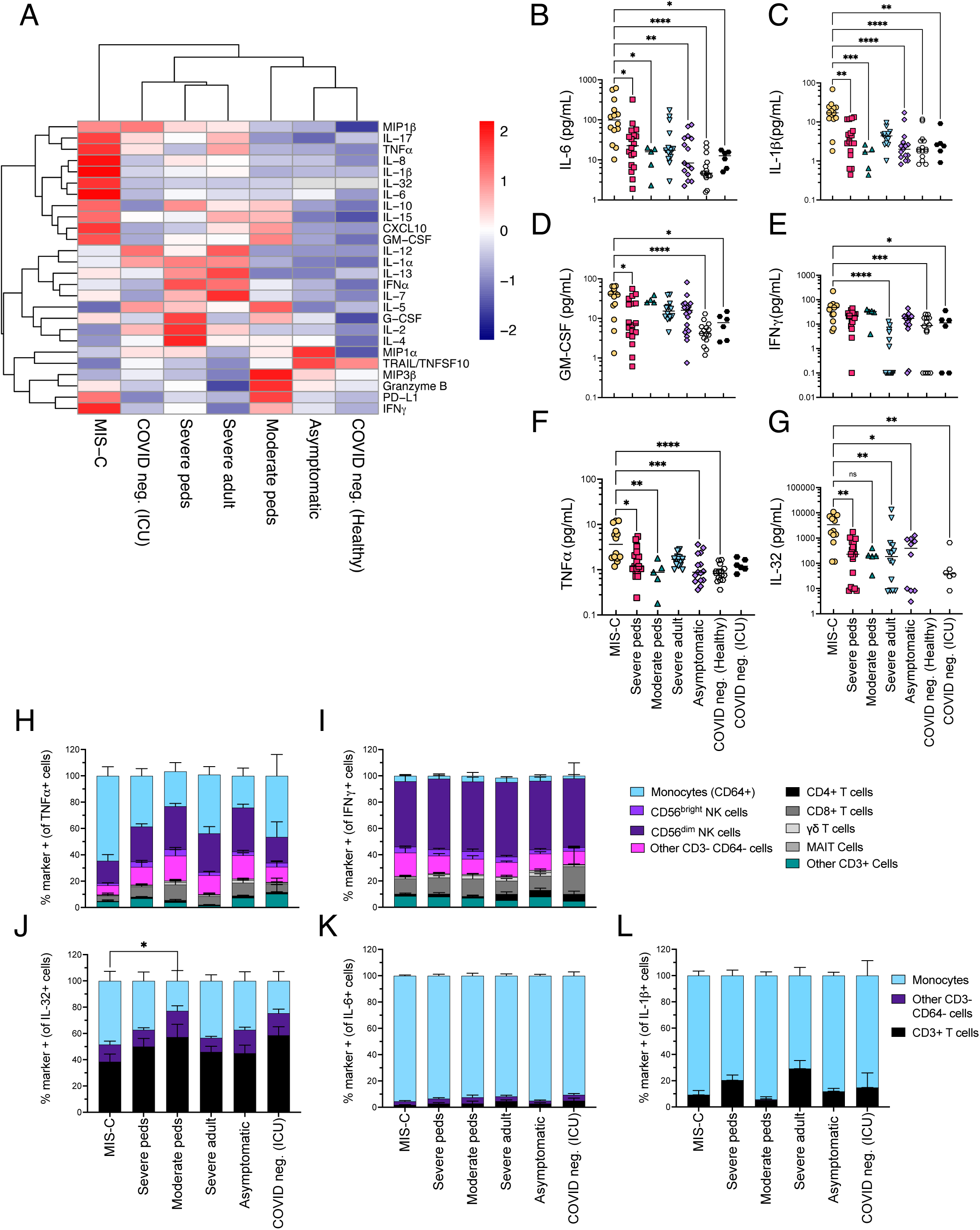
Distinct cytokine and chemokine profiles between MIS-C and acute COVID-19 control groups. **(A)** Profiles of cytokines and chemokines in plasma shown as a heatmap and stratified by group; MIS-C (n=14), severe peds (n = 13), moderate peds (n=5), severe adult (n = 14), asymptomatic (n=15), COVID-19 negative healthy (n=15), COVID-19 negative ICU (n=6). Patients that received convalescent plasma were removed from this analysis. Color intensity of each cell represents the average Z-score of cytokine measures within each group. **(B-G)** Analyte IL-6, Il-1β, GM-CSF, IFNγ, TNFα and IL-32 plasma levels in MIS-C compared to each control group. Statistical analyses were performed using Kruskal-Wallis test with Dunn’s multiple comparison test. Black lines indicate median. *, P <0.05, ** P< 0.01, ***, <0.001, **** P < 0.0001. **(H-L)** Intracellular cytokine production of IFNγ, TNFα, IL-32, IL-6, and IL-1β in an antibody-dependent cellular cytotoxicity/antibody-dependent cellular phagocytosis assay. H-I are from one flow panel and J-L are from a different myeloid cell flow cytometry panel. Statistical analyses were performed using Kruskal-Wallis test with Dunn’s multiple comparison test. *, P <0.05.

We then sought to identify which cytokines produced by Fc receptor-expressed leukocytes may contribute to pathological systemic cytokines. Specifically, we used an antibody-dependent flow cytometry assay to assess intracellular cytokine production across leukocytes. For different cell types we found the proportions of TNFα (Fig.2H), IFNγ (Fig. 2I), IL-32 (Fig.2J), IL-6 (Fig. 2K), and IL-1β (Fig. 2L) in the subsequent functional flow cytometry panels. We found that monocytes produce a high proportion of the pro-inflammatory cytokines (TNFα, IL-32, IL-6, and IL-1β) that were present systemically in children with MIS-C (Fig. 2H, J, K, L). These data suggest that monocytes may be a key driver of pro-inflammatory cytokine production that is seen in MIS-C.

### Increased functionality of monocytes is a distinct feature of MIS-C

Antibody-mediated cellular effector functions correlate with SARS-CoV-2 infection resolution *in vitro* and in animal studies (Chan et al., 2021; Schäfer et al., 2021; Yasui et al., 2014). However, it is not clear whether antibody-mediated cellular functions are protective or pathogenic in MIS-C. To gain additional insights into these cellular functions, we performed an in-depth cellular analysis of innate immune cell phenotypes and function in the different disease groups with *in vitro* functional assays. We first assessed the broad distribution of innate cell types in the blood. We found that MIS-C and severe COVID-19 adult subjects both had decreased total numbers of lymphocytes and monocytes, however, their frequencies were broadly similar. (Fig. S3A-D). Furthermore, the absolute numbers of mucosal associated invariant T cells (MAIT), γδ T cells, CD4+ T cells, and CD56dim NK cells were decreased (Fig. S3C-D).

We next assessed phenotype and function of monocytes. Our analysis included markers of monocyte subsets (CD14, CD16, iNOS, Arginase 1), activation (HLA-DR), Fc receptors (CD16, CD32a, CD32b), cytokine production (IL-1β, IL-6, IL-32, IFNγ, TNFα) and phagocytosis. To study ADCP, we used CFSE- and cholesterol-oligo-Cy5 labeled RBCs as targets (described in Fig. S4A-B and methods) and a hIgG1 anti-CD235a antibody to stimulate the cells to perform ADCP. The myeloid cells or γδ T cells that have phagocytosed cells in this assay are CFSE+ Cy5+ (Fig. 3A).

**Figure 3.**
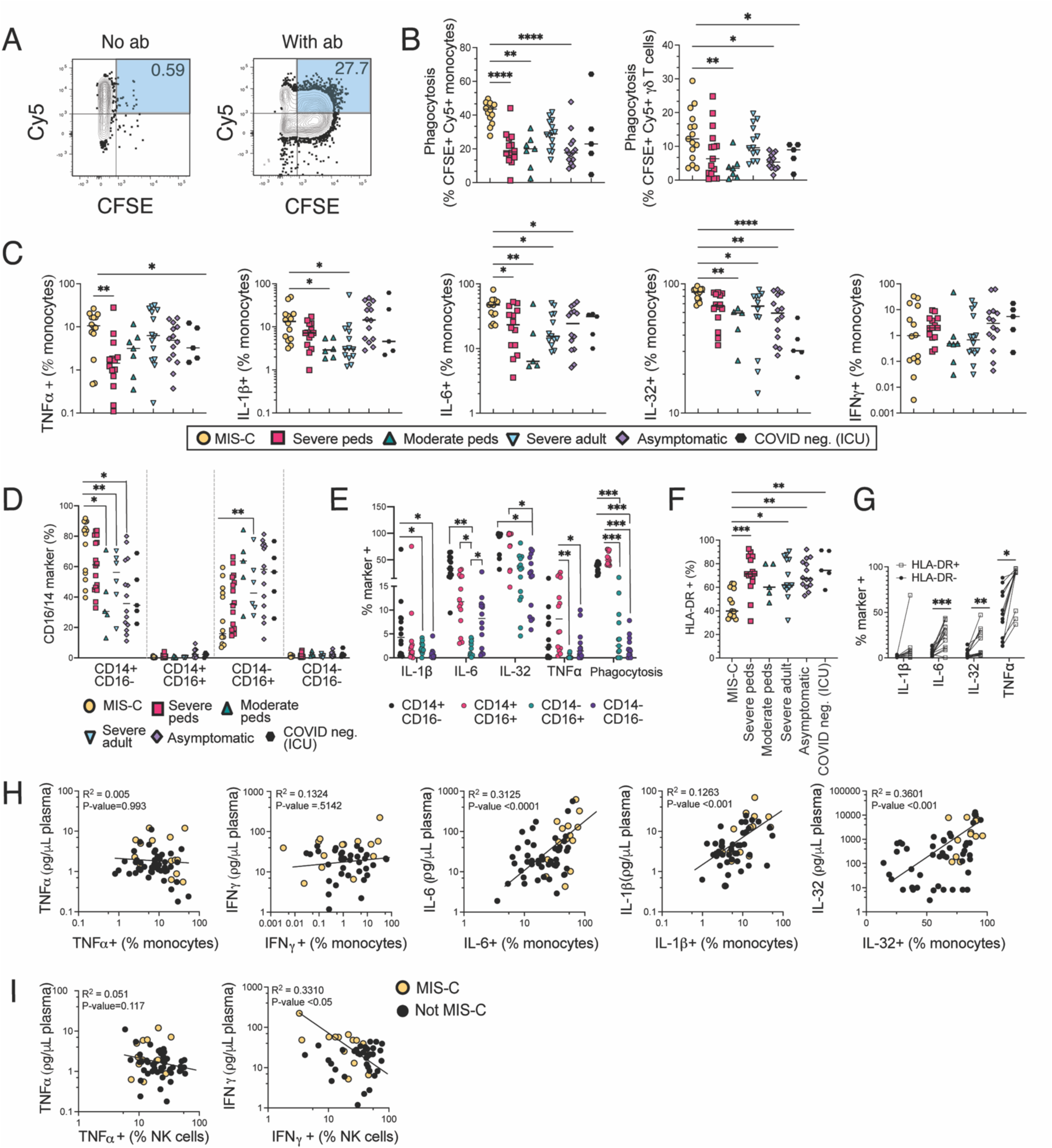
In MIS-C, monocytes and γδ T cells have increased phagocytosis while monocytes have increased cytokine production. **(A)** Combined flow cytometry files from all subjects of CD64+ monocytes showing CFSE by Cy5 on monocytes incubated with RBCs with or without anti-CD235a and labeled with CFSE and cholesterol-oligo-Cy5 probe (n=72). Phagocytosis (%) is defined as percent of cells that are both CFSE+ and Cy5+. **(B-C)** Proportion of monocytes **(B)** and γδ T cells **(C)** performing phagocytosis, as defined by the flow plots in figure 3A, stratified by group. Black lines indicate median. **(C)** Intracellular cytokine staining of cells after incubation with antibody-coated RBC. Proportion of monocytes that produced TNFα, IFNγ, IL-1β, IL-6, and IL-32, stratified by group. Black lines indicate median. (**D)** Proportion of CD16 and CD14 monocyte subsets by group. Black lines indicate median. **(E)** Intracellular cytokine staining and measurement of phagocytosis on monocytes after incubation with antibody-coated RBC stratified based on expression of CD14 and CD16 in MIS-C subjects. **(F)** Proportion of HLA-DR+ cells for monocytes, stratified by group. Black lines indicate median. **(G)** Intracellular cytokine staining and measurement of phagocytosis on monocytes after incubation with antibody-coated RBC stratified based on expression of HLA-DR in MIS-C subjects. For **A-G**, statistical analyses were performed using a Kruskal-Wallis test with Dunn’s multiple comparison test. *, P <0.05, ** P< 0.01. ***P<0.001, **** P < 0.0001. **(H-I)** Proportion of pro-inflammatory cytokines produced by monocytes **(H)** and NK cells **(I)** compared to the plasma levels of respective cytokines by ELISA. R^2^ goodness-of-fit analysis on nonlinear regression line (log transformed) is shown. Yellow dots indicate MIS-C subjects. Black dots indicate non-MIS-C subjects from all other groups for which samples have paired ELISA and flow cytometry data (not all subjects included, n varies depending on flow cytometry test run). P values determined by Pearson correlation. *, P <0.05, ** P< 0.01, ***, <0.001, **** P < 0.0001.

We first assessed the ADCP ability of monocytes and γδ T cells in MIS-C compared to the control groups and found that monocytes and γδ T cells in MIS-C had a greater proportion of cells that phagocytosed target cells (CFSE+ Cy5+) than that of children with acute COVID-19 and asymptomatic COVID-19 infection (Fig. 3B). Adults with severe COVID-19 and children with MIS-C had a similar proportion of cells that phagocytosed target cells. In the same ADCP assay, we can identify which cells produce cytokines in response to being stimulated by an opsonized RBC target. We found that children with MIS-C had a significantly greater proportion of TNFα, IL-1β, IL-6, and IL-32+ monocytes compared to multiple control groups. IFNγ production by monocytes did not differ between groups (Fig. 3C). We also looked at cytokine production from other cell types that have Fc receptors, including mucosal associated invariant T (MAIT) cells and γδτ cells. These cell types can also be stimulated via antibody-mediated mechanisms; however, they comprise a much smaller percentage of lymphocytes in the blood and have not been well studied in MIS-C. TNFα production in MAIT and γδ T cells was similar between groups while IFNγ production was greater in children with MIS-C from MAIT cells compared to other groups (Fig. S3E-F).

We then wanted to characterize the subsets of monocytes producing cytokines in MIS-C. Given that monocytes perform ADCP through engagement of Fc receptors, we first looked at expression of Fc receptors on monocytes including CD64 (FcγRIa), CD32a (FcRγIIa) and CD32b (FcRγIIb) (Anania et al., 2019). We found no differences in expression of these receptors between groups (Fig. S4G). We also assessed if the enhanced ADCP in MIS-C was influenced by Fc receptor genetic variants with different affinity for IgG. To investigate if this could be involved, we assessed a cohort from the Karolinska Institute of MIS-C patients that had whole genome sequencing data. We found no correlation of any Fc receptor tested for the 20 person MIS-C cohort versus published databases matched for ethnicity. Furthermore, the only difference we found was that the G allele in FcγRIA (CD64), the allele that has previously associated with increased CD64 expression and IFNγ production (Wu et al., 2022), was enriched in only 1 out of 20 MIS-C compared to healthy controls (Fig S5 and Table S2). These data suggest that differences in Fc receptor genetics are not driving functional differences in our MIS-C cohort.

We also looked at Arginase-1 and iNOS expression because metabolism via iNOS or Arginase can affect cellular function (Lu et al., 2015; Rodriguez et al., 2017). We found no differences in iNOS or Arginase-1 expression between monocyte groups (Fig. S3G). In addition, human monocytes are divided into three major subsets: classical (CD14+ CD16-), non-classical (CD14dim or neg, CD16+) and intermediate (CD14+ CD16+) (Kapellos et al., 2019). Children with MIS-C had a greater percentage of classical monocytes that were CD14+ CD16- compared to every other group (Fig. 3D). We then investigated the cytokine production by each subset. We found that in children with MIS-C, IL-1β, IL-6, and IL-32 were produced at a higher proportion in classical monocytes (CD14+ CD16-), while TNFα was produced at higher proportions in intermediate (CD14+ CD16+) monocytes. Classical and non-classical monocytes phagocytosed 10-fold more target cells in children with MIS-C. (Fig. 3E). We then investigated HLA-DR expression because it is an activation marker and a decrease in HLA-DR has been found in adult acute severe COVID-19 and sepsis patients as an immunoprotective mechanism (Kim et al., 2010; Leijte et al., 2020). We found that monocytes had significantly decreased expression of HLA-DR in children with MIS-C compared to every other group (Fig. 3F). Those cells that were HLA-DR+ from children with MIS-C were producing more pro-inflammatory cytokines than HLA-DR negative cells, similar to what has been shown in adults with severe COVID-19 infection (Fig. 3G) (Idiz et al., 2022).

We then looked at whether cytokine production by monocytes correlated with the systemic levels of cytokine in the blood to gain insight into which cell subsets may drive disease pathogenesis via cytokine production. The high levels of IL-1β, IL-6, and IL-32 produced by monocytes significantly correlated with corresponding serum levels of these cytokines (Fig. 3H). TNFα and IFNγ serum levels did not correlate with monocyte production of these cytokines. In the ADCP and ADCC assays, we could also assess the production of TNFα and IFNγ by NK cells. Serum levels of TNFα and IFNγ were inversely correlated with NK cell production of these cytokines in the ADCC assay (Fig. 3I).

### Natural killer cells have decreased function and increased expression of exhaustion markers in children with MIS-C

Because we found that NK cell cytokine production was inversely correlated with paired plasma levels, we next wanted to interrogate NK cell phenotype and function across patient groups. In diseases with hyperactive myeloid cells like macrophage activation syndrome (MAS), one commonality is that cellular cytotoxic function is low for genetic or ‘trained’ reasons (Cifaldi et al., 2015; Villanueva et al., 2005). Upon first inspection, we found that both the absolute count and proportion of CD56dim NK cells was decreased in children with MIS-C compared to other acute COVID-19 groups (Fig. S3A-B). We used flow cytometry based *in vitro* assays to understand NK cell phenotype and function in MIS-C (example gating scheme shown in Figure 4A). Overall, CD56dim NK cells from children with MIS-C have increased expression of the exhaustion markers, PD-1, Tim-3, TIGIT and KLRG1, compared to acute pediatric COVID-19 or asymptomatic pediatric subjects. The expression levels of these markers are comparable to that of adult severe COVID-19 subjects (Fig. 4B). There was also a decrease in the inhibitory marker Siglec-7, but no difference in the expression of the inhibitory receptors CTLA-4 and Siglec-9 between groups (Fig. 4B-C).In contrast to what has been observed in adults with COVID-19, children with MIS-C did not have an increase in the adaptive NK cell subset, NKG2C+/CD57+ (Fig. S3H) (Maucourant et al., 2020).

**Figure 4.**
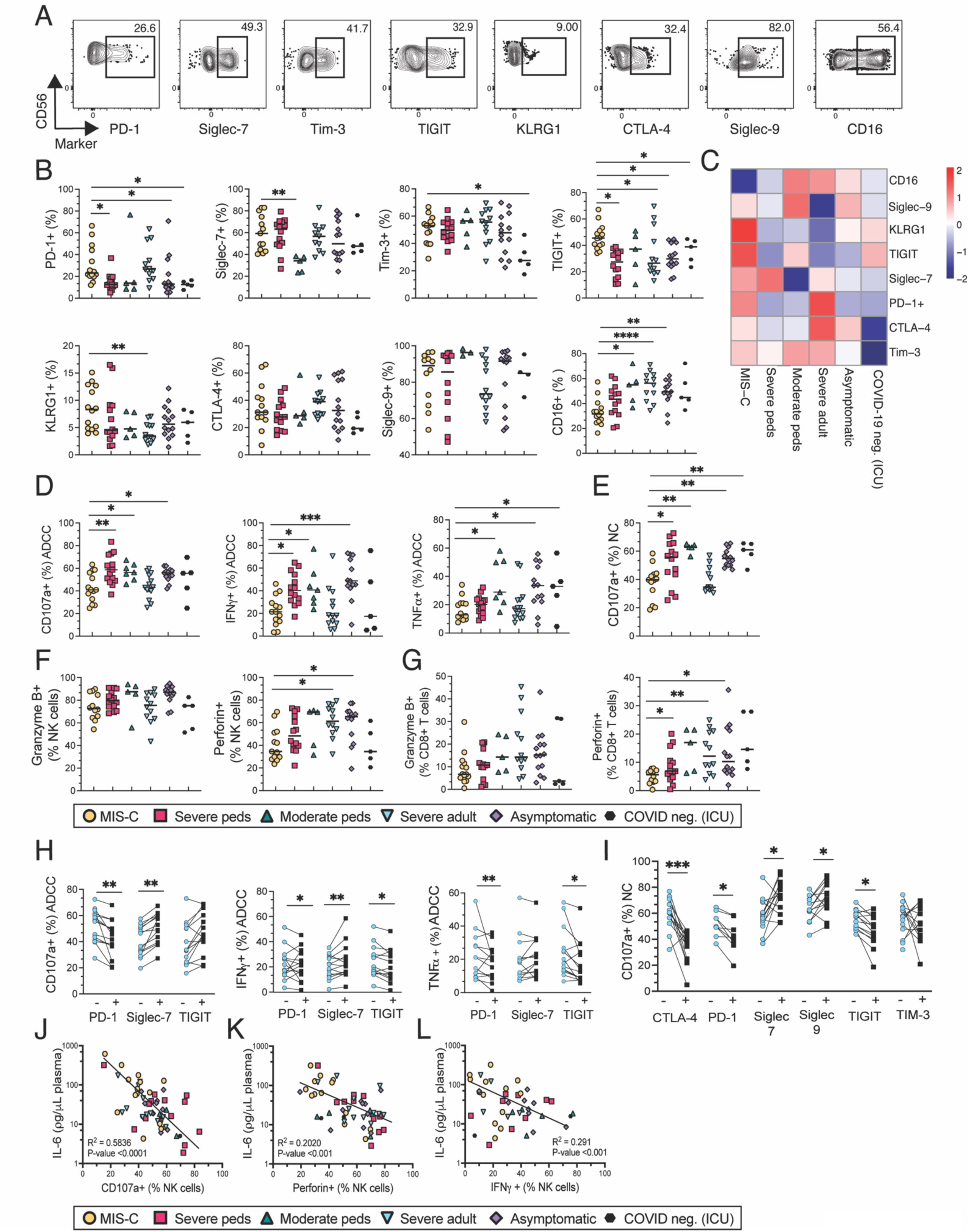
In MIS-C, CD56dim NK cells exhibit an exhausted phenotype and have decreased cytotoxicity. **(A)** Combined flow cytometry plots of all subject for CD56dim NK cells (gated on Singlets, Live Cells, CD3- CD64-) by expression of exhaustion markers (n= 72) **(B)** Proportion of expression of select exhaustion markers on CD56dim NK cells stratified by group; MIS-C (n=14), severe peds (pediatric) (n = 21), moderate peds (pediatric) (n=7), severe adult (n = 14), asymptomatic (n=15) COVID-19 negative ICU (n=6). Black lines indicate median. **(C)** Profiles of exhaustion markers on CD56dim NK cells as shown as a heatmap and stratified by group. Color intensity represents average of marker expression divided the average expression from every subject**. (D-E)** Proportion of CD56dim NK cells degranulated (CD107a) or produced IFNγ, or TNFα in ADCC **(D)** or natural cytotoxicity assay **(E**). Black lines depict median. (**F-G**) Proportion of CD56dim NK cells (**F**) or CD8 T cells (**G**) expressing granzyme B and perforin. For **B-F**, statistical analyses were performed using a Kruskal-Wallis test with Dunn’s multiple comparison test. For **G**, a paired t-test was performed. Black lines depict median. *, P <0.05, ** P< 0.01. ***P<0.001, **** P < 0.0001. **(H-I)** Proportion of CD56dim NK cell degranulation, IFNγ and TNFα production in an ADCC (**H**) or NC (**I**) assay based on expression of exhaustion markers. Statistical analyses were performed using a paired T-test. *, P <0.05, ** P< 0.01, ***, <0.001. **(J-L)** Plasma levels of IL-6 compared to the proportion of degranulating (**J**), perforin positive (**K**) and IFNγ producing (**L**) NK cells. R^2^ goodness-of-fit analysis on log transformed regression line is shown. Statistical testing as done with a Pearson correlation.

Given that NK cells perform ADCC through engagement of its Fc receptor CD16a, we want to look at CD16 expression on NK cells. In an assay that contained no anti-CD235a antibody, NK cells from the MIS-C group had decreased expression of the Fc receptor CD16. It is possible that CD16 was cleaved by matrix metalloproteases, such as ADAM17, *in vivo,* which may contribute to decreased CD16. We also assessed if there might be any influence by Fc receptor genetic variants with different affinity for IgG on NK cell ADCC function. Using genome sequencing data again from the Karolinska Institute, we found no differences in Fc receptor genetics between MIS-C and healthy control that would contribute to differences in ADCC functionality (Table S2), suggesting that any potential differences in NK cell ADCC function in MIS-C would not be due to Fc receptor genetics.

Given the significant proportion of NK cells with exhaustion markers seen in children with MIS-C, we wanted to investigate how well NK cells function via ADCC and natural cytotoxicity (NC). Using flow cytometry based *in vitro* functional assays, we investigated the ability of NK cells to degranulate (CD107a+) and produce pro-inflammatory cytokines, TNFα and IFNγ, in both an ADCC (Fig. 4D) and natural cytotoxicity (NC) assay (Fig. 4E). We found that NK cells from children with MIS-C had significantly decreased amounts of degranulation, TNFα and IFNγ in both ADCC and NC assay compared to other pediatric groups. However, the degranulation and cytokine production of NK cells in children with MIS-C was equivalent to adults with acute severe COVID-19 (Fig. 4D-E). Additionally, NK cells from children with MIS-C had decreased amounts of perforin but no change in Granzyme B expression (Fig. 4F). CD8+ T cells, which are also cytotoxic, also had decreased amounts of perforin, but no change in granzyme B (Fig. 4G).

We then sought to determine how specific subsets of NK cells function. We found that NK cells that were negative for CTLA-4, PD-1, TIGIT and Tim-3 showed increased ADCC and NC functionality compared to NK cells that were positive for those markers (Fig. 4H-I). In contrast, NK cells that were positive for Siglec-7 and Siglec-9 had increased degranulation and cytokine production compared to cells that were negative for those markers for both ADCC and natural cytotoxicity (Fig. 4H-I). This suggests a potential context-dependent hierarchy of exhaustion markers where the increase in some receptors make larger causal impacts on NK cell function and other markers may be less impactful or simply a marker of a subset.

We next wanted to understand what may drive NK cell decreased functionality in MIS-C. Previous work has shown that in COVID-19 adults and children with juvenile idiopathic arthritis that high levels of IL-6 can cause decreased levels of perforin and granzyme A in NK cells and correlate with decreased NK cell function (Cifaldi et al., 2015; Mazzoni et al., 2020). We found that IL-6 levels in the serum (Fig 4J-L), but not IL-1β or IL-32 levels (data not shown), inversely correlated with degranulation, perforin, and IFNγ levels, suggesting that high levels of IL-6 may drive NK cell dysfunction in MIS-C (Fig. 4J-L).

### High-affinity anti-CD16 cellular engagers increases *in vitro* function of NK cells from children with MIS-C and severe adult patients

We next wanted to see if we could improve NK cell function in MIS-C individuals with different FcγRIIIa genotypes through a bi-specific killer engager (BiKE). On one end of the BiKE, we used an anti-CD16 nanobody that binds with a higher affinity to CD16 than the natural Fc portion on IgG to CD16 (Holliger and Hudson, 2005; Vallera et al., 2020; Vallera, 2017). On the other end, we used an anti-CD235a scFv to bind to RBCs that were the proof of principle target (CD16xCD235a BiKE). We assessed NK cell degranulation (CD107a+) when NK cells were stimulated with a CD16xCD235a BiKE compared to a hIgG1 αCD235a mAb. We hypothesized that because the anti-CD16 nanobody is high affinity, this would increase the function of the NK cells for ADCC because of a stronger CD16 signal. We found that degranulation and TNFα was increased for subjects with the homozygous FcγRIIIa SNP 230TT genotype, the genotype with lower affinities to bind to CD16a, to similar proportions as the subjects with the SNP 230TA heterozygous genotype (Fig. 5A; data not shown). Additionally, degranulation was increased only in individuals with the low affinity FcγRIIIa SNP 559TT genotype, the genotype with lower affinities to CD16a but not in heterozygous 559TG or subjects with the high affinity 559GG genotypes (Fig. 5A).

**Figure 5.**
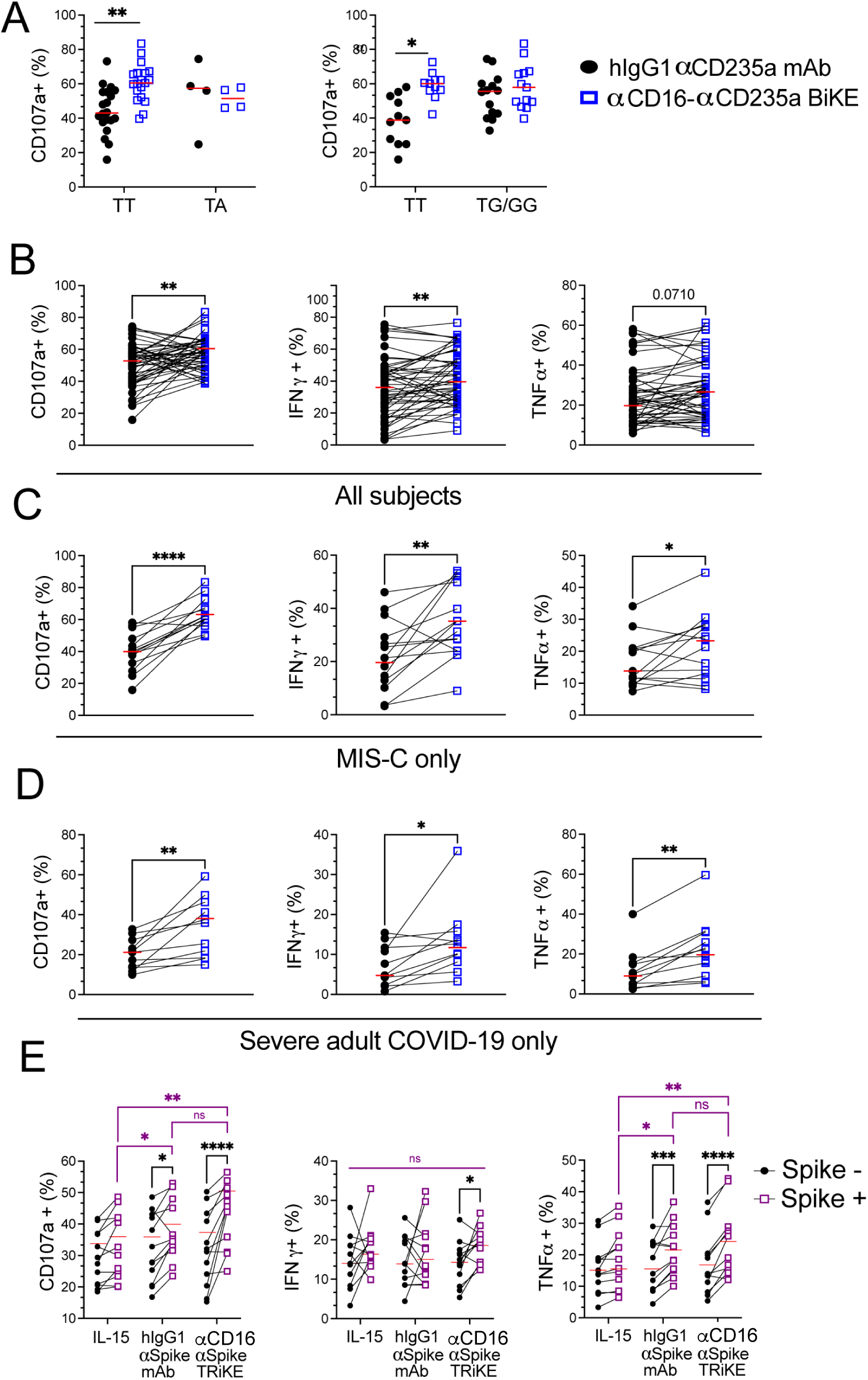
Cellular engagers that target CD16 with high affinity improve NK cell function compared to a human IgG1 antibody. **(A)** Proportion of degranulation in response to an RBC target opsonized with a hIgG1anti-CD235a antibody (black dots) or a αCD16-αCD235a BiKE (open squares) based on FcγR3A SNP (rs10127939 or c.230T>A on the left panel and rs396991 or c.559T>G on the right panel) genotypes. **(B-D)** Proportion of degranulation as measured by CD107a, IFNγ, and TNFα production comparing a hIgG1 αCD235a antibody (black dots) to αCD16-αCD235a BiKE (blue dots) in all subjects **(B)** in MIS-C alone **(C)** or in severe adult subjects alone **(D)**. **(E)** Proportion of NK cells degranulating or producing cytokines as measured by CD107a, IFNγ, and TNFα comparing HEK 293 cells (Spike -) with a HEK 293 Spike-expressing cell line (Spike +) when cultured with IL-15 alone, a hIgG1 α-SARS-CoV-2 Spike mAb (10μg/ml), or αCD16-IL-15-αSARS-CoV-2 Spike TRiKE (10μg/ml). E:T ratio = 3:1. Statistical analyses were performed using a paired t-test. *, P <0.05, ** P< 0.01. Red lines depict median.

We then wanted to see if the CD16xCD235a BiKE, regardless of FcγRIIIa genotype, would improve NK cell function in hypofunctional subjects, such as MIS-C or elderly adults. We found that in response to targets coated by the CD16xCD235a BiKE reagent as opposed to a hIgG1, NK cells degranulated and produced more IFNγ and TNFα in all subjects (Fig. 5B). The NK cell response to the BiKE reagent was greater in MIS-C and acute adults with severe COVID-19 infection compared to the hIgG1 antibody (Fig. 5B-C). Of note, this BiKE did not significantly increase phagocytosis or cytokine production of monocytes, likely because most monocytes in MIS-C are CD16 negative (data not shown).

As a proof of concept, we then developed a tri-specific killer engager (TRiKE) that engages CD16 on one end, the spike protein of SARS-CoV-2 on another end and contains IL-15 to further promote NK cell killing (Arvindam et al., 2021). This reagent differs from the CD16xCD235a BiKE as it is directly targeting the Spike protein on SARS-CoV-2 *in vitro.* We found that the CD16x15xSpike TRiKE significantly increased NK cell degranulation and cytokine production when Spike was present on a target cell surface (Fig. 5 E). This provides proof of concept that an anti-SARS-CoV-2 TRiKE may be useful to enhance NK cell function in the context of infectious diseases.

## Discussion

The immunological mechanisms underlying MIS-C have been difficult to assess because of its relative rarity amongst SARS-CoV-2 infections. As a result, the antibody-mediated immune response in MIS-C is not fully understood. Here, we present antibody and innate immune profiling of children who developed MIS-C compared to other control children and adults with a spectrum of COVID-19 disease severity. We found that children with MIS-C displayed a substantial neutralizing antibody response and a highly inflammatory and distinct cytokine milieu, consistent with what has been shown previously (Rybkina et al., 2023). These observations led us to the hypothesis that even though there are neutralizing antibodies, there is lingering SARS-CoV-2 infection in children with MIS-C because of a dysfunctional antibody-mediated cellular response. Using *ex vivo* flow cytometry assays, we investigated antibody-mediated effector functions across the COVID-19 disease spectrum. We found that children with MIS-C have aberrant activation of monocytes, which, although important for clearing virus, may also contribute to the inflammation leading to the pathology of MIS-C. In contrast, we found that NK cells have decreased functionality, an exhaustion marker signature, and that more systemic IL-6 correlates with reduced function of NK cells. This suggests that NK cell dysfunction potentially contributes (by being absent) to MIS-C pathogenesis. Also, the dysfunction in the ADCC response in children with MIS-C and severe adults can be rescued and target SARS-CoV-2 with high affinity anti-CD16 BiKE and TRiKE molecules respectively. To our knowledge, this is the first study to report on antibody-dependent cellular functions for children with MIS-C as compared to a spectrum of COVID-19 control groups. Together, our results help identify specific cells that may be causing the pathogenic inflammation in children with MIS-C.

Children with MIS-C exhibit a unique cytokine and chemokine profile in the plasma, with higher levels of the pro-inflammatory cytokines, consistent with previous reports of cytokines in MIS-C serum (Butters et al., 2023; Consiglio et al., 2020; Gurlevik et al., 2022; Hoste et al., 2022; Rybkina et al., 2023; Sacco et al., 2022). Interestingly, we found high amounts of IL-32 in MIS-C, suggesting that IL-32 may be a new biomarker of MIS-C. Relevant to this study, IL-32 can induce the differentiation of monocytes and promote the production of TNFα and IL-1β. Previous reports have shown that IL-32, IFNγ, and IL-6 were the best at distinguishing severe forms of COVID-19 in adults (Bergantini et al., 2022; Khawar et al., 2016; Kim et al., 2005). However, our IL-32 levels in children with MIS-C were even higher as compared to the severe adult COVID-19 group. Therefore, further work is needed on IL-32 to see if it would be an additional marker of MIS-C and to see if its affects are causing pathology. Children with MIS-C also had much higher levels of IL-10 than any other group. We postulate this amount of IL-10 is being produced to combat the high levels of inflammation but is not enough to offset the onslaught of pro-inflammatory cytokines that are found in the blood.

An important mechanistic next step is knowing which cells produce the systemic cytokines that are elevated in MIS-C; this will enable cell-specific therapies to be included in treatment approaches. It was unknown which Fc receptor+ cells produce cytokines in response to antibody stimulation in children with MIS-C. In ADCP assays, we found that monocytes from children with MIS-C constituted a greater proportion of IL-1β, IL-6, TNFα, and IL-32 positive cells. Furthermore, monocytes and γδ T cells from MIS-C exhibited greater levels of phagocytosis compared to those from adult and pediatric individuals with acute COVID-19 in the same assay. In this way, monocytes are functioning as a “double-edged sword” in MIS-C – on the one side, SARS-CoV-2 specific antibody-mediated phagocytosis can eliminate free virus or cells that are making SARS-CoV-2. While on the other side, production of pro-inflammatory cytokines can contribute to disease pathology. We propose that even though monocytes may decrease viral load, the cytokines produced by monocytes, like IL-6, negatively impact the response of other cell types that are beneficial for viral clearance, like NK cells.

For other diseases with hyperactive myeloid cells, such as MAS, an underlying correlate can be a lack of activity in cytotoxic cells such as NK cells. Little is known about NK function in children with MIS-C and no work has yet studied whether the strength of the NK cell response correlates with disease severity. Furthermore, a severe gap in knowledge is the lack of research on the ADCC responses of NK cells from individuals with COVID-19 infection (Lee and Blish, 2023). When we examined NK cell natural cytotoxicity (NC) and ADCC function in MIS-C, we found that NK cells in MIS-C had decreased degranulation and cytokine production in NC and ADCC. Furthermore, NK cells in children with MIS-C had decreased perforin; this is likely attributed to NK cell training, as opposed to a primary perforin defect based on previous studies in severe COVID-19 in adults (Kundura et al., 2024). One hypothesis for why the NK cells may be hypofunctional was that the pro-inflammatory milieu drove them to exhaustion. We found some evidence for this as NK cells in MIS-C had significantly greater proportions of an exhaustion marker signature (PD-1+, TIGIT+, Tim-3+, KLRG1+). Details on the combined or individual functions of these exhaustion markers in NK cells still need to be better understood. Additionally, previous studies have shown that IL-6 downregulated the NK cell and CD8+ T cell cytotoxic response through the STAT3 pathway (Cifaldi et al., 2015; Dhar et al., 2021; Wu et al., 2019). We found that higher plasma IL-6 levels correlated with reduced NK cell function and perforin levels, suggesting that high levels of IL-6 produced by monocytes may partially explain the decreased NK cell function in MIS-C. Our data suggest that IL-6 may lead to NK cell dysfunction/exhaustion, and that the exhaustion markers we found that correlated with reduced function could be a signature of IL-6 induced exhaustion. Further work will need to be done to understand if reduced activity of NK cells and/or other cytolytic cells is a symptom or cause of the hyperinflammatory MIS-C disease. If lack of activity by NK cells is causal, identifying the most critical primary functions of NK cells, such as killing virally infected cells or killing stressed inflammatory cells, would be important for treatment of MIS-C and key to understanding their role in protection.

Considering the NK cell ADCC dysfunction in children with MIS-C, we wanted to determine if we could increase their NK cell ADCC function. One way to target an antibody-like response to a particular cell type is to use Bi-specific Killer Engagers (BiKEs) or Tri-specific Killer Engagers (TRiKEs). BiKEs and TRiKEs that target CD16 on NK cells have been utilized in multiple different cancers (Myers and Miller, 2021; Pawlowski et al., 2023). Therefore, we hypothesized that the BiKE reagent would generate a stronger response than a human IgG1 antibody. We found that this approach worked to align the NK cells that were hypofunctional in MIS-C and adults with severe COVID-19 symptoms to the maximal response of the other groups in the study. Importantly, CD16 has two isoforms, CD16a and CD16b. CD16a is expressed on NK cells and CD16b is expressed on neutrophils. Development of a CD16a specific nanobody would be more ideal to target NK cells. We then tested a TRiKE that targeted CD16 on NK cells and SARS-CoV-2 Spike protein and found significantly increased NK cell degranulation when Spike was on the surface of the target cell *in vitro*. Our proof of principle data suggest that a CD16-Spike TRiKE could be used in SARS-CoV-2 to improve the antibody-mediated response of NK cells. Therefore, NK-cell targeted therapies may be a useful strategy for helping to control SARS-CoV-2 viral clearance, while not adding to the pathological cytokine response from monocytes.

In summary, homeostasis is essential to clearance of virus without causing immunopathology. In the case of the MIS-C antibody-mediated response, while it is likely that multiple pathogenic mechanisms contribute to MIS-C, our data suggest an imbalance of the antibody-mediated cellular response as a mechanistic correlate of this rare, but life-threatening, outcome of SARS-CoV-2 infection in children. Furthermore, if MIS-C is indeed driven by dysregulated innate immunity responsible for ineffective viral and/or pathological cell clearance, then targeted therapies to diminish monocyte cytokine production and boost NK cell responses may be effective in treating MIS-C or similar infectious disease phenotypes.

## Materials and Methods

### Human subjects

From May 2020 to May 2021, we prospectively enrolled MIS-C and control pediatric patients (0-21y.o.) from Children’s Hospitals and Clinics of Minnesota (Minneapolis and St. Paul, MN campuses). To classify a patient as MIS-C, we followed the MIS-C case definition by the World Health Organization and the Center for Disease Control; <21 years old, fever of >38 °C for 24 hrs, laboratory evidence of inflammation, hospital admission, multi-organ involvement, no alternative plausible diagnosis, and positive SARS-CoV-2 serology (Vogel et al., 2021). All samples were collected from subjects during routine phlebotomy performed for clinical purposes. The patients and family members provided informed consent or assent if written consent was provided by a minor’s parent in accordance with the 1975 (as revised in 2000/2008) Helsinki Declaration for enrollment in research protocols that were approved by the Institutional Review Board (IRB) of the University of Minnesota and Children’s Hospital of Minnesota. In acute COVID-19 patients, severity was determined based on symptoms, respiratory support requirements, and severity of organ injury (Gallo Marin et al., 2021; Health, 2024; Jonat et al., 2021). Of note, individuals that were in the COVID-19 positive asymptomatic pediatric controls group came to the hospital for some reason other than SARS-CoV-2 and were enrolled in this study because they were incidentally found to be SARS-CoV-2 positive by nuclei acid testing.

Additionally, we obtained cryopreserved adult specimens from the University of Minnesota (UMN) COVID-19 Biorepository and plasma waste specimens from MIS-C patient samples as part of the UMN Biorepository from December 2020 to May 2021. We also obtained COVID-19 negative pediatric healthy control and COVID-19 positive asymptomatic pediatric controls samples from the Development of Immunity after SARS-CoV-2 Exposure and Recovery (DISCOVER) study at the Indiana University School of Medicine. These samples were collected at Riley Hospital for Children (Indianapolis, IN) and the details regarding sample collection can be found in previous publications (Khaitan et al., 2022). We also utilized sequencing data from 20 additional MIS-C samples collected at the Karolinska Institute. These samples were only used to assess Fc receptor genetic associations with MIS-C relative to published data sets. All samples were collected pre-vaccination against SARS-CoV-2. Samples for MIS-C and acute pediatric groups were obtained within 72 hours (hrs) of hospitalization. Samples from adults were collected between 3 and 14 days into their hospitalization. IRB approval was obtained at all institutions.

We pre-specified exclusion criteria and therefore excluded subjects from both the studies and the assays performed. In all groups, individuals with documented immunodeficiency or malignancy were excluded. Additionally, samples were excluded from assays if insufficient numbers of PBMCs could be recovered or if >50% of the cells were dead based on flow cytometry staining. For the detection of SARS-CoV-2 specific antibodies, we excluded samples that had received convalescent plasma as this result could be confounded by the antigen specific antibodies given to the patient in the convalescent plasma. Therefore, not every assay has the same number of subjects. Characteristics for subjects are found in Tables 1 and 2.

### Primary Cell Purification and Freezing

For the blood samples, Ficoll centrifugation was used to isolate PBMCs, plasma and red blood cell pellets (containing large leukocytes and RBCs). The top portion was taken as plasma and placed in a −80° C freezer. The buffy coat layer in the middle was taken for PBMCs. Lastly, the RBC pellet layer at the bottom was stored at −80° C for later DNA purification. Once isolated, PBMCs were counted and resuspended in 500 μl of RMPI-1640 + 50% FBS and transferred to a cryovial. When all tubes were ready to freeze, 500 μl of FBS +15% DMSO was added to the cryovials with cells (7.5% final DMSO concentration). The cells were then immediately transferred to a pre-cooled Mr. Frosty and placed in −80° C freezer. After 24 hrs, the cells were transferred to a vapor-phase liquid nitrogen freezer for long-term storage. For the DISCOVER samples, PBMCs were separated from venous blood by polysucrose and sodium diatrizoate density gradient centrifugation (Sigma-Aldrich, St. Louis, MO) and cryopreserved in liquid nitrogen after resuspension heat-inactivated FBS and 10% DMSO as described previously (Khaitan et al., 2022).

The primary red blood cells (RBCs) used as antibody target cells in assays were obtained and purified from deidentified adult blood donors (Memorial Blood Center). RBCs were purified from whole blood by leukocyte reduction filtration (Fenwal Inc, RS-2000) and resuspended at 50% hematocrit in RPMI-1640 and 25 mM HEPES, L-Glutamine and 50 mg/l Hypoxanthine.

### Cell line maintenance and culture

The K562 cell line was maintained between 1 × 10^5^ and 1.5 × 10^6^ cells/ml in RPMI 1640 with 10% heat inactivated FBS and 1 mg/ml gentamicin (termed RP10) and incubated at 37°C with 5% CO_2_.

The HEK 293 T and a SARS-CoV-2 Spike expressing HEK 293 cell line (293 SARS2) were maintained between 1 × 10^5^ and 1.5 × 10^6^ cells/ml in Dulbecco’s Modified Eagle Medium with 10% heat inactivated FBS and 100mU Pen-Strep. The 293-SARS2 cell line was also incubated with 100μg/mL Normocin and 10μg/mL blasticidin to maintain Spike expression. The cells were incubated at 37°C with 5% CO_2_.

### Measurement of serum cytokine and chemokine levels

Samples were tested by the Cytokine Reference Laboratory at the University of Minnesota (CLIA’88 licensed #24D0931212). For bead-based cytokine detection, samples were analyzed for 25 human specific analytes using a multiplex platform. Samples were assayed according to manufacturer’s instructions. The samples were read on a Luminex-based instrument (Bioplex 200). Samples were run in duplicate, and values were interpolated from 5-parameter fitted standard curves created using recombinant proteins.

For quantification of human IL-32, samples were analyzed using a quantitative sandwich enzyme-linked immunosorbent assay (ELISA) platform. Samples were assayed according to manufacturer’s instructions. The absorbance was measured on the microtiter plate reader (EPOCH). Samples were tested in duplicate, and values were interpolated from a LOG-LOG fitted standard curve.

### Quantification of anti-SARS-CoV-2 antibodies

An indirect ELISA was used for quantification of anti-SARS-CoV-2 antibodies from plasma. For Spike proteins, 96 well polystyrene high-binding ELISA plates were coated overnight with 66.7nM of the coating proteins (Spike 1, Spike 2) in 50 mM Na_2_CO_3_, pH 9.6, with bovine serum albumin (BSA) as a negative control. For the membrane and envelope peptides, biotinylated peptides were coated onto streptavidin plates at room temperature (RT) for 2 hrs with 66.7nM of each peptide or an anti-human IgG-Biotin as a positive control at 66.7nM. Peptide sequences are found in Table S1. Plates were then washed 5 times with PBS+0.05% Tween-20 (PBST) before being loaded with 3-fold serially diluted plasma samples (1:200-1:1,312,200) and incubated for 1.5 hrs at RT on an orbital shaker at 60 RPM and then for 30 minutes (min) at RT without shaking. Plates were subsequently washed 5 times with PBST before incubation with a horseradish peroxidase (HRP)-conjugated goat anti-human IgG at a 1:50,000 dilution overnight at 4°C. The plates were finally washed 5 times with PBST. HRP activity was quantitated by incubating with o-phenylenediamine substrate for 30 minutes, and then the reaction was stopped by addition of 2% oxalic acid. Finally, the plates were read by absorbance at 492 nm on a 96-well plate reader (Tecan Infinite) to assess SARS-CoV-2-specific IgG titers (OD_492_). Values were plotted on dilution curves, and area under the curve (AUC) was calculated using GraphPad Prism software version 10. The mean OD_492_ values of the blank control was used to set a background value in the analysis.

### Plaque reduction neutralization test

Plaque reduction neutralization test (PRNT) procedures were performed in a biosafety level 3 facility approved by the Institutional Biosafety Committee at the University of Minnesota. Plasma samples were heat inactivated at 56°C for 30 min, then serially diluted two-fold in Dulbecco’s Modified Eagle Medium (DMEM). They were then mixed with 200 plaque forming units (PFU) of SARS-CoV-2 isolate 2019-nCoV/USA-WA1/2020 (NR-52281, BEI Resources, NIAID, NIH) in a total volume of 250 µl. The plasma-virus mixtures were incubated for 1h at 37°C and 100 µl of plasma-virus mixtures were transferred to confluent Vero E6 cell monolayers in duplicate wells of 24-well tissue culture plates. Plates were incubated for 1hr with intermittent mixing at 37°C / 5% CO_2_. After 1 hr, 500 µl of overlay medium (DMEM + 1.6% microcrystalline cellulose + 2% FBS) was added to each well and incubated for 48 hrs at 37°C with 5% CO_2_. Then the cells were fixed in a 4% paraformaldehyde solution and incubated for 30 min at RT. The cells were then washed two times with phosphate buffer saline (PBS), and then stained with 0.1% crystal violet to visualize viral plaques. A virus preparation without addition of plasma was used as the negative control. 50% plaque reduction neutralization titer (PRNT_50_) was determined as the reciprocal of the highest dilution of plasma that showed ≥50% reduction in the number of SARS-CoV-2 plaques.

### Generation of α-CD235a and α-Spike protein recombinant antibody reagents

The sequence of the variable heavy chain (VH) and variable light chain (VL) domains of the human glycophorin A (CD235a) binding murine hybridoma-derived antibody was first described as clone 10F7(Bigbee et al., 1983; Way, 2017). It was expressed recombinantly on a human IgG1/IgKappa scaffold or as a bispecific killer engager (BiKE) containing the α-CD235a as a single-chain variable fragment (scFv) fused to a nanobody specific for human CD16 (Felices et al., 2020; Vallera et al., 2020). The anti-CD235a hIgG1 reagent was cloned, expressed, and purified following the previously described recombinant antibody production process (Hicks et al., 2022). VH and VL gene fragments for Gibson assembly were codon optimized and synthesized by Twist Bioscience. The amino acid (AA) sequences of the neutralizing (CCL6.29) SARS-CoV-2 binding variable heavy (VH) and variable light (VL) antibody domains were first described in Rogers et al. (Rogers et al., 2020). It was expressed recombinantly as a tri-specific killer engager (TRiKE) DNA shuffling and ligation techniques connected a camelid anti-CD16 complementarity determining region splice into a humanized scaffold; via linter regions to human IL-15; and then on to an the anti-SARS-CoV-2 scFv, as previously described (Felices et al., 2020).

The gene fragment encoding the CD16xCD235a BiKE and the CD16x15xSpike TRiKE with a N-terminal signal peptide and a C-terminal 10x-Histidine tag was synthesized by Integrated DNA Technologies and cloned into a pMC.EF1α-MCS-SV40polyA Parental Minicircle Cloning Vector (System Biosciences) via Gibson assembly using previously described methods (Vallera et al., 2020). The Expi293 expressed CD16xCD235a BiKE and CD16x15xSpike TRiKE was purified using nickel-affinity chromatography with HisTrap excel resin (Cytiva).

### Antibody-dependent cellular cytotoxicity assay with RBCs as targets

RBCs were incubated with either monoclonal anti-CD235a hIgG1 or CD16xCD235a BiKE. Then they were combined with PBMCs at a PBMC:RBC ratio of 1:1 in RP10 with 1µg/ml brefeldin A, 1µg/ml monensin, and 1:200 anti-CD107a. As a negative control, PBMCs were also incubated with RBC alone (no antibody). The cells were then incubated at 37°C for 4 hrs. After 4 hrs, the cells were centrifuged, washed, and were prepared for flow cytometry.

### Antibody-dependent cellular cytotoxicity assay with HEK 293 T and 293 SARS2 cell lines as targets

HEK 293 T and 293 SARS2 cells were stained with a 2 µM solution of carboxyfluorescein succinimidyl ester (CFSE) at 37°C for 30 min in PBS. The cells were then washed 3 times with RP10 and counted. The HEK293 T cells were incubated with either monoclonal anti-Spike hIgG1 or CD16x15xSpike TRiKE at 10 µg. Then they were combined with NK cells at a E:T ratio of 3:1 in RP10 with 1µg/ml brefeldin A, 1µg/ml monensin, and 1:200 anti-CD107a. As a negative control, NK cells were also incubated with IL-15 alone. The cells were then incubated at 37°C for 5 hrs. After 5 hrs, the cells were centrifuged, washed with FACS buffer, and were prepared for flow cytometry.

### Antibody-dependent phagocytosis assays

RBCs were stained with a 2 µM solution of carboxyfluorescein succinimidyl ester (CFSE) at 37°C for 30 min in PBS. The cells were then washed 3 times with RP10 and resuspended in 200 µL of RP10 in a 96-well plate overnight in a 37°C incubator. The next morning, CFSE-labeled RBCs were counted and labeled with 300 nM of a Cy5-oligo-cholesterol probe (Table S1 for sequence), and the RBCs were incubated at RT for 30 min in the dark. The probe was modified from the original design as previously described (Ana-Sosa-Batiz et al., 2014). The RBCs were then centrifuged and washed 3 times with RP10. RBCs were then opsonized with either monoclonal anti-CD235a hIgG1 or CD16xCD235a BiKE and combined with PBMCs at a ratio of 1:1 in RP10 with 1µg/ml brefeldin A and 1µg/ml monensin. As a negative control, PBMCs were also incubated with RBCs alone (no antibody or BiKE). The cells were then incubated at 37°C for 4 hrs. After 4 hrs, the cells were centrifuged, washed, and resuspend in 500 nM of a fluorescence quencher attached to a reverse complement oligo to quench Cy5 signal on non-phagocytosed RBCs (Table S1 for sequences) at RT for 20 min in the dark. The cells were centrifuged and washed three times with RP10 and then were prepared for flow cytometry.

### Natural cytotoxicity assay

K562 cells and PBMCs were combined at a PBMC: K562 ratio of 1:1 in RP10 + 1µg/ml brefeldin A, 1µg/ml monensin and 1:200 anti-CD107a. The cells were then incubated at 37°C for 4 hrs in a V bottom 96 well plate. After 4 hrs, the cells were centrifuged, washed, and prepared for flow cytometry (see below).

### Flow cytometry

Once the ADCC, ADCP, and NC assays were complete, cells were resuspended in a master mix made of PBS and fixable viability dye, then incubated at RT in the dark for 20 min in a 96-well V bottom plate. The cells were washed with PBS. The cells were then resuspended in PBS, 2% FBS, 2 mM EDTA (FACS buffer) with surface stain antibody master mix, then incubated at RT in the dark for 20 min. The cells were washed with FACS buffer and incubated in 2% formaldehyde at 37°C for 10 min. After incubation, the cells were washed and resuspended in 0.04% Triton X-100 at RT in the dark for exactly 7 minutes, then washed in FACS buffer with 2% BSA and stained in FACS buffer with 2% BSA and internal stain mastermix. The cells were then incubated at RT in the dark for 30 min. After incubation, the cells were washed with FACS buffer and resuspended in FACS buffer before acquiring data on a LSRFortessa flow cytometer (BD Biosciences) or CytoFLEX flow cytometer (Beckman Coulter). Data was analyzed with FlowJo 10.8.0 software. The antibodies used are listed in Table S1.

### Genotyping of *FCGR3A* SNPs used for functional flow cytometry analysis

*FCGR* family member genes were generated through duplication and divergence during evolution (Qiu et al., 1990). We used a modified published *FCGR* SNP TaqMan assay in which *FCGR* gene-specific PCR fragments were used as templates instead of genomic DNA for TaqMan assays (Wu et al., 2022; Wu et al., 2014). Briefly, genomic DNA was isolated from the red blood cell pellets of donors using the Wizard Genomic DNA Purification Kit and the genomic DNA fragments containing functional SNPs of *FCGR3A* were amplified using the gene specific primers as described previously (Wu et al., 2022; Wu et al., 2014). All TaqMan assays were designed using the Software Primer Express v3.0 (Applied Biosystems Inc.). TaqMan genotyping assays were carried out according to the standard protocol on an ABI 7500 Real-Time PCR System machine using Genotyping Master Mix (Applied Biosystems).

### Genotyping of *FCGR* SNPs used for associations with MIS-C

Sequencing data from 20 children with MIS-C were also obtained from a cohort at the Karolinska Institute in Sweden. Genomic DNA from the Swedish cohort was extracted from whole blood using Puregene Blood Core Kit C (Qiagen). Illumina whole-genome sequencing, mapping, and read processing and annotation was performed according to the MIP Rare disease pipeline utilized by the Genomic Medicine Center Karolinska Rare Diseases (GMCK-RD) (Stranneheim et al., 2021). The ethnicity of these sequences was assessed using principle component analysis (PCA) to be able to compare if the *FCGR* SNPs of a particular ethnicity in the MIS-C cohort from the Karolinska Institute was significantly different than that of the 1000 Genomes Projects (1kGP) (Auton et al., 2015). For generating the PCA plot, patient and 1kGP vcfs were merged. Variant pruning and PCA was performed using plink2 and principal components were visualized in R with the ggplot2 package. SNP genotype data was extracted for patients and selected controls using vcftools (Danecek et al., 2011).

### Assessing sequence variations in SARS-CoV-2 genes

Individual FASTA files for the Spike, membrane, and envelope protein sequences were obtained from Global Initiative on Sharing All Influenza Data (GISAID) (Shu and McCauley, 2017). Sequences are available upon request. Multiple sequence alignments were performed using the online multiple sequence aligner MAFFT v7.4 (Multiple Alignment program for amino acid or nucleotide sequences) using the first U.S case as a reference (GenBank: MN985325.1) and Percent Accepted Mutation (PAM) value of 1 (Holshue et al., 2020). We filtered out any sequences with >5% ambiguous nucleotide fraction. Default parameters were used except for a default offset value of 0.1. Following alignment, the individual open reading frames for the Spike, envelope, and membrane proteins were visualized using Jalview (Waterhouse et al., 2009). The percent amino acid changes relative to the reference were calculated using NCBI’s Nucleotide Sequence Alignment Viewer (Sayers et al., 2022). Protein topology and extra-virion portions were predicted using MEGAv11 and confirmed with published experimental evidence in available cases (Schoeman and Fielding, 2019; Thomas, 2020; Troyano-Hernáez et al., 2021).

### Heatmap

For the heatmaps, values below the lower limit of quantification (LLOQ) were set to equal the lowest non-LLOQ value and were summarized for each category (e.g., MIS-C) by taking the median value for the individuals in that category. Summarized data were mean-centered and scaled for each marker. Each value had the mean of all values for that marker subtracted from it, and then each centered value was divided by the standard deviation for that marker. The heatmap was drawn using R (v.4.3.0) package heatmap (v.1.0.12) and clustering was done using Euclidean distance and complete linkage (2023; Kolde, 2018).

### Statistical analysis

Excel 16.54 (Microsoft) and Prism 9.3.1 (GraphPad Software) were used for data transformation, generation of figures, and statistical analyses. Illustrator 25.4.1 (Adobe) was used to draft figures for publication. Sample sizes, independent experiments run, statistical tests, and significance thresholds are listed in each figure legend. * = p < 0.05, ** = p < 0.01, *** = p < 0.001, **** = p < 0.0001.

## Data and code availability

All reagents used in this study are listed in Table S3 and S4. Further information and requests for data or reagents should be directed to the corresponding author, Geoffrey T Hart.

## Acknowledgements

This work was supported by the Department of Medicine, Department of Pharmacology, the Department of Veterinary Medicine, the Clinical and Translational Science Institute, the Department of Pediatrics, Division of Pediatric Critical Care Medicine at the University of Minnesota. Also supported by the National Institutes of Health grants ULTR1002494, KL2TR002392, T32AI007313-34A1, Children’s Hospital of Minnesota, and the Children’s Health Discovery Fund. The authors declare no competing financial interests.

Special thanks to the staff at the University of Minnesota flow cytometry core and Minnesota Supercomputing Institute. Thanks to the innovative work by Dr. Martin Felices and Todd Lenvik the CD16 BiKE and TRiKE collaboration. We thank the University of Minnesota, BSL-3 program staff for their support of the high containment research laboratories used for these studies. The SARS-CoV-2 isolate hCoV-19/USA-WA1/2020 NR-52281 was deposited by the Centers for Disease Control and Prevention and obtained through BEI Resources, NIAID, NIH. We would like to thank the National Cancer Institute (NCI) Biological Resources Branch for providing human IL-15. Thanks to Dr. Tyler Bold, Dr. Peter Southern, Dr. Luca Schifanella, and Dr. Christopher Tignanelli for their work to generate and maintain the University of Minnesota (UMN) COVID-19 Biorepository. We are grateful to the parents and patients who generously provided samples for our studies. Multiple figures created with BioRender.com.

## Author Contributions

Conceptualization: GTH.; Methodology: JKD, VDK, DH, KE, RAK, AJ, CB, JSM, BKT, JW, MC, MP, GTH.; Investigation: JKD, JAS, VDK, AK, LH, SE, AJ, LEC, CB, CS, BKT, CMH, JW, MES, G.T.H.; Validation: JKD, CMH, GTH; Formal Analysis: JKD, VDK, AK, AJ, LEC, CMH, BKT, GTH; Resources: DH, YS, PB, AKhaitan, YTB, CCJ, AO, MES; Data Curation: JKD, GTH; Writing – Original Draft: JKD, GTH; Writing – Review & Editing: JKD, VDK, DH, LH, AJ, KE, CB, LEC, YTB, CS, CCJ, JSM, YTB, JW, CCJ, AO, MCC, AKhaitan; Visualization: JKD, GTH; Supervision: GTH; Funding Acquisition: JKD, AO, MES, MP, GTH; Project Administration: GTH.

## Declaration of interests

The authors declare no competing interests related to the content of this work.

Acronyms: antibody dependent cellular cytotoxicity (ADCC), antibody dependent cellular phagocytosis (ADCP), bi-specific killer engager (BiKE), Coronavirus disease of 2019 (COVID-19), multisystem inflammatory syndrome in children (MIS-C), natural killer (NK) cells, PBMC (Peripheral Blood Mononuclear Cells), tri-specific killer engager (TRiKE)

## Acronyms

ADCC: antibody dependent cellular cytotoxicity
ADCP: antibody dependent cellular phagocytosis
BiKE: bi-specific killer engager
COVID-19: Coronavirus disease of 2019
MIS-C: multisystem inflammatory syndrome in children
NK (cells): natural killer
PBMC: Peripheral Blood Mononuclear Cells
TRiKE: tri-specific killer engager

## References

2023.R: A Language and environment for statistical computing. In R Foundation for Statistical Computing, Vienna, Austria.

Adeniji, O.S., L.B. Giron, M. Purwar, N.F. Zilberstein, A.J. Kulkarni, M.W. Shaikh, R.A. Balk, J.N. Moy, C.B. Forsyth, Q. Liu, H. Dweep, A. Kossenkov, D.B. Weiner, A. Keshavarzian, A. Landay, and M. Abdel-Mohsen. 2021. COVID-19 Severity Is Associated with Differential Antibody Fc-Mediated Innate Immune Functions. mBio 12:

Akkoyun, E.B., Z. Most, H. Katragadda, A. Yu, L. Nassi, N. Oakman, S. Ginsburg, and M. Maamari. 2023. Impact of anakinra use on clinical outcomes in children with moderate or severe multisystem inflammatory syndrome in children: a propensity score matched retrospective cohort study. Pediatr Rheumatol Online J 21:141.

Ana-Sosa-Batiz, F., A.P.R. Johnston, H. Liu, R.J. Center, S. Rerks-Ngarm, P. Pitisuttithum, S. Nitayaphan, J. Kaewkungwal, J.H. Kim, N.L. Michael, A.D. Kelleher, I. Stratov, S.J. Kent, and M. Kramski. 2014. HIV-specific antibody-dependent phagocytosis matures during HIV infection. Immunology & Cell Biology 92:679–687.

Anania, J.C., A.M. Chenoweth, B.D. Wines, and P.M. Hogarth. 2019. The Human FcγRII (CD32) Family of Leukocyte FcR in Health and Disease. Frontiers in Immunology 10:

Andersson, U. 2021. Hyperinflammation: On the pathogenesis and treatment of macrophage activation syndrome. Acta Paediatrica 110:2717–2722.

Arvindam, U.S., P.M.M. van Hauten, D. Schirm, N. Schaap, W. Hobo, B.R. Blazar, D.A. Vallera, H. Dolstra, M. Felices, and J.S. Miller. 2021. A trispecific killer engager molecule against CLEC12A effectively induces NK-cell mediated killing of AML cells. Leukemia 35:1586–1596.

Auton, A., G.R. Abecasis, D.M. Altshuler, R.M. Durbin, G.R. Abecasis, D.R. Bentley, A. Chakravarti, A.G. Clark, P. Donnelly, E.E. Eichler, P. Flicek, S.B. Gabriel, R.A. Gibbs, E.D. Green, M.E. Hurles, B.M. Knoppers, J.O. Korbel, E.S. Lander, C. Lee, H. Lehrach, E.R. Mardis, G.T. Marth, G.A. McVean, D.A. Nickerson, J.P. Schmidt, S.T. Sherry, J. Wang, R.K. Wilson, R.A. Gibbs, E. Boerwinkle, H. Doddapaneni, Y. Han, V. Korchina, C. Kovar, S. Lee, D. Muzny, J.G. Reid, Y. Zhu, J. Wang, Y. Chang, Q. Feng, X. Fang, X. Guo, M. Jian, H. Jiang, X. Jin, T. Lan, G. Li, J. Li, Y. Li, S. Liu, X. Liu, Y. Lu, X. Ma, M. Tang, B. Wang, G. Wang, H. Wu, R. Wu, X. Xu, Y. Yin, D. Zhang, W. Zhang, J. Zhao, M. Zhao, X. Zheng, E.S. Lander, D.M. Altshuler, S.B. Gabriel, N. Gupta, N. Gharani, L.H. Toji, N.P. Gerry, A.M. Resch, P. Flicek, J. Barker, L. Clarke, L. Gil, S.E. Hunt, G. Kelman, E. Kulesha, R. Leinonen, W.M. McLaren, R. Radhakrishnan, A. Roa, D. Smirnov, R.E. Smith, I. Streeter, A. Thormann, I. Toneva, B. Vaughan, X. Zheng-Bradley, D.R. Bentley, R. Grocock, S. Humphray, T. James, Z. Kingsbury, H. Lehrach, R. Sudbrak, M.W. Albrecht, V.S. Amstislavskiy, T.A. Borodina, M. Lienhard, F. Mertes, M. Sultan, B. Timmermann, M.-L. Yaspo, E.R. Mardis, R.K. Wilson, L. Fulton, R. Fulton, S.T. Sherry, V. Ananiev, Z. Belaia, D. Beloslyudtsev, N. Bouk, C. Chen, D. Church, R. Cohen, C. Cook, J. Garner, T. Hefferon, M. Kimelman, C. Liu, J. Lopez, P. Meric, C. O’Sullivan, Y. Ostapchuk, L. Phan, S. Ponomarov, V. Schneider, E. Shekhtman, K. Sirotkin, D. Slotta, H. Zhang, G.A. McVean, R.M. Durbin, S. Balasubramaniam, J. Burton, P. Danecek, T.M. Keane, A. Kolb-Kokocinski, S. McCarthy, J. Stalker, M. Quail, J.P. Schmidt, C.J. Davies, J. Gollub, T. Webster, B. Wong, Y. Zhan, A. Auton, C.L. Campbell, Y. Kong, A. Marcketta, R.A. Gibbs, F. Yu, L. Antunes, M. Bainbridge, D. Muzny, A. Sabo, Z. Huang, J. Wang, L.J.M. Coin, L. Fang, X. Guo, X. Jin, G. Li, Q. Li, Y. Li, Z. Li, H. Lin, B. Liu, R. Luo, H. Shao, Y. Xie, C. Ye, C. Yu, F. Zhang, H. Zheng, H. Zhu, C. Alkan, E. Dal, F. Kahveci, G.T. Marth, E.P. Garrison, D. Kural, W.-P. Lee, W. Fung Leong, M. Stromberg, A.N. Ward, J. Wu, M. Zhang, M.J. Daly, M.A. DePristo, R.E. Handsaker, D.M. Altshuler, E. Banks, G. Bhatia, G. del Angel, S.B. Gabriel, G. Genovese, N. Gupta, H. Li, S. Kashin, E.S. Lander, S.A. McCarroll, J.C. Nemesh, R.E. Poplin, S.C. Yoon, J. Lihm, V. Makarov, A.G. Clark, S. Gottipati, A. Keinan, J.L. Rodriguez-Flores, J.O. Korbel, T. Rausch, M.H. Fritz, A.M. Stütz, P. Flicek, K. Beal, L. Clarke, A. Datta, J. Herrero, W.M. McLaren, G.R.S. Ritchie, R.E. Smith, D. Zerbino, X. Zheng-Bradley, P.C. Sabeti, I. Shlyakhter, S.F. Schaffner, J. Vitti, D.N. Cooper, E.V. Ball, P.D. Stenson, D.R. Bentley, B. Barnes, M. Bauer, R. Keira Cheetham, A. Cox, M. Eberle, S. Humphray, S. Kahn, L. Murray, J. Peden, R. Shaw, E.E. Kenny, M.A. Batzer, M.K. Konkel, J.A. Walker, D.G. MacArthur, M. Lek, R. Sudbrak, V.S. Amstislavskiy, R. Herwig, E.R. Mardis, L. Ding, D.C. Koboldt, D. Larson, K. Ye, S. Gravel, C. The Genomes Project, a. Corresponding, c. Steering, g. Production, M. Baylor College of, B.G.I. Shenzhen, M.I.T. Broad Institute of, Harvard, R. Coriell Institute for Medical, E.B.I. European Molecular Biology Laboratory, Illumina, G. Max Planck Institute for Molecular, U. McDonnell Genome Institute at Washington, U.S.N.I.o. Health, O. University of, I. Wellcome Trust Sanger, g. Analysis, Affymetrix, M. Albert Einstein College of, U. Bilkent, C. Boston, L. Cold Spring Harbor, U. Cornell, L. European Molecular Biology, U. Harvard, D. Human Gene Mutation, S. Icahn School of Medicine at Mount, U. Louisiana State, H. Massachusetts General, U. McGill, and N.I.H. National Eye Institute. 2015. A global reference for human genetic variation. Nature 526:68–74.

Bahnan, W., S. Wrighton, M. Sundwall, A. Bläckberg, O. Larsson, U. Höglund, H. Khakzad, M. Godzwon, M. Walle, E. Elder, A.S. Strand, L. Happonen, O. André, J.K. Ahnlide, T. Hellmark, V. Wendel-Hansen, R.P. Wallin, J. Malmstöm, L. Malmström, M. Ohlin, M. Rasmussen, and P. Nordenfelt. 2021. Spike-Dependent Opsonization Indicates Both Dose-Dependent Inhibition of Phagocytosis and That Non-Neutralizing Antibodies Can Confer Protection to SARS-CoV-2. Front Immunol 12:808932.

Bergantini, L., M. d’Alessandro, P. Cameli, A. Otranto, S. Luzzi, F. Bianchi, and E. Bargagli. 2022. Cytokine profiles in the detection of severe lung involvement in hospitalized patients with COVID-19: The IL-8/IL-32 axis. Cytokine 151:155804.

Bi, J., and Z. Tian. 2017. NK Cell Exhaustion. Front Immunol 8:760.

Bigbee, W.L., M. Vanderlaan, S.S. Fong, and R.H. Jensen. 1983. Monoclonal antibodies specific for the M- and N-forms of human glycophorin A. Mol Immunol 20:1353–1362.

Burbelo, P.D., R. Castagnoli, C. Shimizu, O.M. Delmonte, K. Dobbs, V. Discepolo, A. Lo Vecchio, A. Guarino, F. Licciardi, U. Ramenghi, E. Rey-Jurado, C. Vial, G.L. Marseglia, A. Licari, D. Montagna, C. Rossi, G.A. Montealegre Sanchez, K. Barron, B.M. Warner, J.A. Chiorini, Y. Espinosa, L. Noguera, L. Dropulic, M. Truong, D. Gerstbacher, S. Mató, J. Kanegaye, A.H. Tremoulet, E.M. Eisenstein, H.C. Su, L. Imberti, M.C. Poli, J.C. Burns, L.D. Notarangelo, and J.I. Cohen. 2022. Autoantibodies Against Proteins Previously Associated With Autoimmunity in Adult and Pediatric Patients With COVID-19 and Children With MIS-C. Front Immunol 13:841126.

Butters, C., N. Benede, T. Moyo-Gwete, S.I. Richardson, U. Rohlwink, M. Shey, F. Ayres, N.P. Manamela, Z. Makhado, S.R. Balla, M. Madzivhandila, A. Ngomti, R. Baguma, H. Facey-Thomas, T.F. Spracklen, J. Day, H. van der Ross Debbie, C. Riou, W.A. Burgers, C. Scott, L. Zuhlke, P.L. Moore, R.S. Keeton, and K. Webb. 2023. Comparing the immune abnormalities in MIS-C to healthy children and those with inflammatory disease reveals distinct inflammatory cytokine production and a monofunctional T cell response. Clin Immunol 109877.

Carter, M.J., M. Fish, A. Jennings, K.J. Doores, P. Wellman, J. Seow, S. Acors, C. Graham, E. Timms, J. Kenny, S. Neil, M.H. Malim, S.M. Tibby, and M. Shankar-Hari. 2020. Peripheral immunophenotypes in children with multisystem inflammatory syndrome associated with SARS-CoV-2 infection. Nat Med 26:1701–1707.

Chakraborty, S., J.C. Gonzalez, B.L. Sievers, V. Mallajosyula, S. Chakraborty, M. Dubey, U. Ashraf, B.Y. Cheng, N. Kathale, K.Q.T. Tran, C. Scallan, A. Sinnott, A. Cassidy, S.T. Chen, T. Gelbart, F. Gao, Y. Golan, X. Ji, S. Kim-Schulze, M. Prahl, S.L. Gaw, S. Gnjatic, T.U. Marron, M. Merad, P.S. Arunachalam, S.D. Boyd, M.M. Davis, M. Holubar, C. Khosla, H.T. Maecker, Y. Maldonado, E.D. Mellins, K.C. Nadeau, B. Pulendran, U. Singh, A. Subramanian, P.J. Utz, R. Sherwood, S. Zhang, P. Jagannathan, G.S. Tan, and T.T. Wang. 2022. Early non-neutralizing, afucosylated antibody responses are associated with COVID-19 severity. Sci Transl Med 14:eabm7853.

Chan, C.E.Z., S.G.K. Seah, H. Chye, S. Massey, M. Torres, A.P.C. Lim, S.K.K. Wong, J.J.Y. Neo, P.S. Wong, J.H. Lim, G.S.L. Loh, D. Wang, J.D. Boyd-Kirkup, S. Guan, D. Thakkar, G.H. Teo, K. Purushotorman, P.E. Hutchinson, B.E. Young, J.G. Low, P.A. MacAry, H. Hentze, V.S. Prativadibhayankara, K. Ethirajulu, J.E. Comer, C.K. Tseng, A.D.T. Barrett, P.J. Ingram, T. Brasel, and B.J. Hanson. 2021. The Fc-mediated effector functions of a potent SARS-CoV-2 neutralizing antibody, SC31, isolated from an early convalescent COVID-19 patient, are essential for the optimal therapeutic efficacy of the antibody. PLoS One 16:e0253487.

Cifaldi, L., G. Prencipe, I. Caiello, C. Bracaglia, F. Locatelli, F. De Benedetti, and R. Strippoli. 2015. Inhibition of natural killer cell cytotoxicity by interleukin-6: implications for the pathogenesis of macrophage activation syndrome. Arthritis Rheumatol 67:3037–3046.

Consiglio, C.R., N. Cotugno, F. Sardh, C. Pou, D. Amodio, L. Rodriguez, Z. Tan, S. Zicari, A. Ruggiero, G.R. Pascucci, V. Santilli, T. Campbell, Y. Bryceson, D. Eriksson, J. Wang, A. Marchesi, T. Lakshmikanth, A. Campana, A. Villani, P. Rossi, N. Landegren, P. Palma, and P. Brodin. 2020. The Immunology of Multisystem Inflammatory Syndrome in Children with COVID-19. Cell 183:968–981.e967.

Crayne, C.B., S. Albeituni, K.E. Nichols, and R.Q. Cron. 2019. The Immunology of Macrophage Activation Syndrome. Frontiers in Immunology 10:

Cui, X., Z. Zhao, T. Zhang, W. Guo, W. Guo, J. Zheng, J. Zhang, C. Dong, R. Na, L. Zheng, W. Li, Z. Liu, J. Ma, J. Wang, S. He, Y. Xu, P. Si, Y. Shen, and C. Cai. 2021. A systematic review and meta-analysis of children with coronavirus disease 2019 (COVID-19). J Med Virol 93:1057–1069.

Danecek, P., A. Auton, G. Abecasis, C.A. Albers, E. Banks, M.A. DePristo, R.E. Handsaker, G. Lunter, G.T. Marth, S.T. Sherry, G. McVean, R. Durbin, and G.P.A. Group. 2011. The variant call format and VCFtools. Bioinformatics 27:2156–2158.

Dhar, S.K., V. K. S. Damodar, S. Gujar, and M. Das. 2021. IL-6 and IL-10 as predictors of disease severity in COVID-19 patients: results from meta-analysis and regression. Heliyon 7:e06155.

Diorio, C., R. Shraim, L.A. Vella, J.R. Giles, A.E. Baxter, D.A. Oldridge, S.W. Canna, S.E. Henrickson, K.O. McNerney, F. Balamuth, C. Burudpakdee, J. Lee, T. Leng, A. Farrel, M.P. Lambert, K.E. Sullivan, E.J. Wherry, D.T. Teachey, H. Bassiri, and E.M. Behrens. 2021. Proteomic profiling of MIS-C patients indicates heterogeneity relating to interferon gamma dysregulation and vascular endothelial dysfunction. Nat Commun 12:7222.

Farooqi, K.M., A. Chan, R.J. Weller, J. Mi, P. Jiang, E. Abrahams, A. Ferris, U.S. Krishnan, N. Pasumarti, S. Suh, A.M. Shah, M.P. DiLorenzo, P. Zachariah, J.D. Milner, E.B. Rosenzweig, M. Gorelik, and B.R. Anderson. 2021. Longitudinal Outcomes for Multisystem Inflammatory Syndrome in Children. Pediatrics 148:

Feldstein, L.R., E.B. Rose, S.M. Horwitz, J.P. Collins, M.M. Newhams, M.B.F. Son, J.W. Newburger, L.C. Kleinman, S.M. Heidemann, A.A. Martin, A.R. Singh, S. Li, K.M. Tarquinio, P. Jaggi, M.E. Oster, S.P. Zackai, J. Gillen, A.J. Ratner, R.F. Walsh, J.C. Fitzgerald, M.A. Keenaghan, H. Alharash, S. Doymaz, K.N. Clouser, J.S. Giuliano, Jr., A. Gupta, R.M. Parker, A.B. Maddux, V. Havalad, S. Ramsingh, H. Bukulmez, T.T. Bradford, L.S. Smith, M.W. Tenforde, C.L. Carroll, B.J. Riggs, S.J. Gertz, A. Daube, A. Lansell, A. Coronado Munoz, C.V. Hobbs, K.L. Marohn, N.B. Halasa, M.M. Patel, and A.G. Randolph. 2020. Multisystem Inflammatory Syndrome in U.S. Children and Adolescents. N Engl J Med 383:334–346.

Felices, M., T.R. Lenvik, B. Kodal, A.J. Lenvik, P. Hinderlie, L.E. Bendzick, D.K. Schirm, M.F. Kaminski, R.T. McElmurry, M.A. Geller, C.E. Eckfeldt, D.A. Vallera, and J.S. Miller. 2020. Potent Cytolytic Activity and Specific IL15 Delivery in a Second-Generation Trispecific Killer Engager. Cancer Immunol Res 8:1139–1149.

Filippatos, F., E.B. Tatsi, and A. Michos. 2023. Immunology of Multisystem Inflammatory Syndrome after COVID-19 in Children: A Review of the Current Evidence. Int J Mol Sci 24:

Gallo Marin, B., G. Aghagoli, K. Lavine, L. Yang, E.J. Siff, S.S. Chiang, T.P. Salazar-Mather, L. Dumenco, M.C. Savaria, S.N. Aung, T. Flanigan, and I.C. Michelow. 2021. Predictors of COVID-19 severity: A literature review. Rev Med Virol 31:1–10.

Grom, A.A., J. Villanueva, S. Lee, E.A. Goldmuntz, M.H. Passo, and A. Filipovich. 2003. Natural killer cell dysfunction in patients with systemic-onset juvenile rheumatoid arthritis and macrophage activation syndrome. J Pediatr 142:292–296.

Gurlevik, S.L., Y. Ozsurekci, E. Sağ, P. Derin Oygar, S. Kesici, Ü.K. Akca, M.K. Cuceoglu, O. Basaran, S. Göncü, J. Karakaya, A.B. Cengiz, and S. Özen. 2022. The difference of the inflammatory milieu in MIS-C and severe COVID-19. Pediatric Research

Health, N.I.o. 2024. COVID-19 Treatment Guidelines Panel. Coronavirus Disease 2019 (COVID-19) Treatment Guidelines. In.

Hicks, D., C. Baehr, P. Silva-Ortiz, A. Khaimraj, D. Luengas, F.A. Hamid, and M. Pravetoni. 2022. Advancing humanized monoclonal antibody for counteracting fentanyl toxicity towards clinical development. Hum Vaccin Immunother 18:2122507.

Holliger, P., and P.J. Hudson. 2005. Engineered antibody fragments and the rise of single domains. Nat Biotechnol 23:1126–1136.

Hollis, N.D., W. Li, M.E. Van Dyke, G.J. Njie, H.M. Scobie, E.M. Parker, A. Penman-Aguilar, and K.E.N. Clarke. 2021. Racial and Ethnic Disparities in Incidence of SARS-CoV-2 Infection, 22 US States and DC, January 1-October 1, 2020. Emerg Infect Dis 27:1477–1481.

Holshue, M.L., C. DeBolt, S. Lindquist, K.H. Lofy, J. Wiesman, H. Bruce, C. Spitters, K. Ericson, S. Wilkerson, A. Tural, G. Diaz, A. Cohn, L. Fox, A. Patel, S.I. Gerber, L. Kim, S. Tong, X. Lu, S. Lindstrom, M.A. Pallansch, W.C. Weldon, H.M. Biggs, T.M. Uyeki, and S.K. Pillai. 2020. First Case of 2019 Novel Coronavirus in the United States. N Engl J Med 382:929–936.

Hoste, L., L. Roels, L. Naesens, V. Bosteels, S. Vanhee, S. Dupont, C. Bosteels, R. Browaeys, N. Vandamme, K. Verstaen, J. Roels, K.F.A. Van Damme, B. Maes, E. De Leeuw, J. Declercq, H. Aegerter, L. Seys, U. Smole, S. De Prijck, M. Vanheerswynghels, K. Claes, V. Debacker, G. Van Isterdael, L. Backers, K.B.M. Claes, P. Bastard, E. Jouanguy, S.Y. Zhang, G. Mets, J. Dehoorne, K. Vandekerckhove, P. Schelstraete, J. Willems, P. Stordeur, S. Janssens, R. Beyaert, Y. Saeys, J.L. Casanova, B.N. Lambrecht, F. Haerynck, and S.J. Tavernier. 2022. TIM3+ TRBV11-2 T cells and IFNγ signature in patrolling monocytes and CD16+ NK cells delineate MIS-C. J Exp Med 219:

Hoste, L., R. Van Paemel, and F. Haerynck. 2021. Multisystem inflammatory syndrome in children related to COVID-19: a systematic review. Eur J Pediatr 180:2019–2034.

Idiz, U.O., T.T. Yurttas, S. Degirmencioglu, B. Orhan, E. Erdogan, H. Sevik, and M.M. Sevinc. 2022. Immunophenotyping of lymphocytes and monocytes and the status of cytokines in the clinical course of Covid-19 patients. J Med Virol

Jia, H., H. Yang, H. Xiong, and K.Q. Luo. 2023. NK cell exhaustion in the tumor microenvironment. Front Immunol 14:1303605.

Jonat, B., M. Gorelik, A. Boneparth, A.S. Geneslaw, P. Zachariah, A. Shah, L. Broglie, J. Duran, K.D. Morel, M. Zorrilla, L. Svoboda, C. Johnson, J. Cheng, M.C. Garzon, W.G. Silver, K. Gross Margolis, C. Neunert, I. Lytrivi, J. Milner, S.G. Kernie, and E.W. Cheung. 2021. Multisystem Inflammatory Syndrome in Children Associated With Coronavirus Disease 2019 in a Children’s Hospital in New York City: Patient Characteristics and an Institutional Protocol for Evaluation, Management, and Follow-Up. Pediatr Crit Care Med 22:e178–e191.

Junqueira, C., Â. Crespo, S. Ranjbar, L.B. de Lacerda, M. Lewandrowski, J. Ingber, B. Parry, S. Ravid, S. Clark, M.R. Schrimpf, F. Ho, C. Beakes, J. Margolin, N. Russell, K. Kays, J. Boucau, U. Das Adhikari, S.M. Vora, V. Leger, L. Gehrke, L.A. Henderson, E. Janssen, D. Kwon, C. Sander, J. Abraham, M.B. Goldberg, H. Wu, G. Mehta, S. Bell, A.E. Goldfeld, M.R. Filbin, and J. Lieberman. 2022. FcγR-mediated SARS-CoV-2 infection of monocytes activates inflammation. Nature 606:576–584.

Kapellos, T.S., L. Bonaguro, I. Gemünd, N. Reusch, A. Saglam, E.R. Hinkley, and J.L. Schultze. 2019. Human Monocyte Subsets and Phenotypes in Major Chronic Inflammatory Diseases. Front Immunol 10:2035.

Khaitan, A., D. Datta, C. Bond, M. Goings, K. Co, E.O. Odhiambo, L. Miller, L. Zhang, S. Beasley, J. Poorbaugh, and C.C. John. 2022. Level and Duration of IgG and Neutralizing Antibodies to SARS-CoV-2 in Children with Symptomatic or Asymptomatic SARS-CoV-2 Infection. Immunohorizons 6:408–415.

Khawar, M.B., M.H. Abbasi, and N. Sheikh. 2016. IL-32: A Novel Pluripotent Inflammatory Interleukin, towards Gastric Inflammation, Gastric Cancer, and Chronic Rhino Sinusitis. Mediators Inflamm 2016:8413768.

Kim, O.Y., A. Monsel, M. Bertrand, P. Coriat, J.M. Cavaillon, and M. Adib-Conquy. 2010. Differential down-regulation of HLA-DR on monocyte subpopulations during systemic inflammation. Crit Care 14:R61.

Kim, S.H., S.Y. Han, T. Azam, D.Y. Yoon, and C.A. Dinarello. 2005. Interleukin-32: a cytokine and inducer of TNFalpha. Immunity 22:131–142.

Knoll, R., J.L. Schultze, and J. Schulte-Schrepping. 2021. Monocytes and Macrophages in COVID-19. Frontiers in Immunology 12:

Kolde, R. 2018. pheatmap: Pretty Heatmaps. R package version 1.0.12. In.

Kundura, L., R. Cezar, E. Ballongue, S. André, M. Michel, C. Mettling, C. Lozano, T. Vincent, L. Muller, J.Y. Lefrant, C. Roger, P.G. Claret, S. Duvnjak, P. Loubet, A. Sotto, T.A. Tran, J. Estaquier, and P. Corbeau. 2024. Low Percentage of Perforin-Expressing NK Cells during Severe SARS-CoV-2 Infection: Consumption Rather than Primary Deficiency. J Immunol 212:1105–1112.

Lazova, S., Y. Dimitrova, D. Hristova, I. Tzotcheva, and T. Velikova. 2022. Cellular, Antibody and Cytokine Pathways in Children with Acute SARS-CoV-2 Infection and MIS-C-Can We Match the Puzzle? Antibodies (Basel) 11:

Lee, M.J., and C.A. Blish. 2023. Defining the role of natural killer cells in COVID-19. Nature Immunology

Lee, P.Y., M. Day-Lewis, L.A. Henderson, K.G. Friedman, J. Lo, J.E. Roberts, M.S. Lo, C.D. Platt, J. Chou, K.J. Hoyt, A.L. Baker, T.M. Banzon, M.H. Chang, E. Cohen, S.D. de Ferranti, A. Dionne, S. Habiballah, O. Halyabar, J.S. Hausmann, M.M. Hazen, E. Janssen, E. Meidan, R.W. Nelson, A.A. Nguyen, R.P. Sundel, F. Dedeoglu, P.A. Nigrovic, J.W. Newburger, and M.B.F. Son. 2020. Distinct clinical and immunological features of SARS-CoV-2-induced multisystem inflammatory syndrome in children. J Clin Invest 130:5942–5950.

Leijte, G.P., T. Rimmelé, M. Kox, N. Bruse, C. Monard, M. Gossez, G. Monneret, P. Pickkers, and F. Venet. 2020. Monocytic HLA-DR expression kinetics in septic shock patients with different pathogens, sites of infection and adverse outcomes. Critical Care 24:110.

Lu, G., R. Zhang, S. Geng, L. Peng, P. Jayaraman, C. Chen, F. Xu, J. Yang, Q. Li, H. Zheng, K. Shen, J. Wang, X. Liu, W. Wang, Z. Zheng, C.F. Qi, C. Si, J.C. He, K. Liu, S.A. Lira, A.G. Sikora, L. Li, and H. Xiong. 2015. Myeloid cell-derived inducible nitric oxide synthase suppresses M1 macrophage polarization. Nat Commun 6:6676.

Matchett, W.E., V. Joag, J.M. Stolley, F.K. Shepherd, C.F. Quarnstrom, C.K. Mickelson, S. Wijeyesinghe, A.G. Soerens, S. Becker, J.M. Thiede, E. Weyu, S.D. O’Flanagan, J.A. Walter, M.N. Vu, V.D. Menachery, T.D. Bold, V. Vezys, M.K. Jenkins, R.A. Langlois, and D. Masopust. 2021. Cutting Edge: Nucleocapsid Vaccine Elicits Spike-Independent SARS-CoV-2 Protective Immunity. J Immunol 207:376–379.

Maucourant, C., I. Filipovic, A. Ponzetta, S. Aleman, M. Cornillet, L. Hertwig, B. Strunz, A. Lentini, B. Reinius, D. Brownlie, A. Cuapio, E.H. Ask, R.M. Hull, A. Haroun-Izquierdo, M. Schaffer, J. Klingström, E. Folkesson, M. Buggert, J.K. Sandberg, L.I. Eriksson, O. Rooyackers, H.G. Ljunggren, K.J. Malmberg, J. Michaëlsson, N. Marquardt, Q. Hammer, K. Strålin, and N.K. Björkström. 2020. Natural killer cell immunotypes related to COVID-19 disease severity. Sci Immunol 5:

Mazzoni, A., L. Salvati, L. Maggi, M. Capone, A. Vanni, M. Spinicci, J. Mencarini, R. Caporale, B. Peruzzi, A. Antonelli, M. Trotta, L. Zammarchi, L. Ciani, L. Gori, C. Lazzeri, A. Matucci, A. Vultaggio, O. Rossi, F. Almerigogna, P. Parronchi, P. Fontanari, F. Lavorini, A. Peris, G.M. Rossolini, A. Bartoloni, S. Romagnani, F. Liotta, F. Annunziato, and L. Cosmi. 2020. Impaired immune cell cytotoxicity in severe COVID-19 is IL-6 dependent. J Clin Invest 130:4694–4703.

Moebius, J., R. Guha, M. Peterson, K. Abdi, J. Skinner, S. Li, G. Arora, B. Traore, S. Rajagopalan, E.O. Long, and P.D. Crompton. 2020. PD-1 Expression on NK Cells in Malaria-Exposed Individuals Is Associated with Diminished Natural Cytotoxicity and Enhanced Antibody-Dependent Cellular Cytotoxicity. Infect Immun 88:

Myers, J.A., and J.S. Miller. 2021. Exploring the NK cell platform for cancer immunotherapy. Nat Rev Clin Oncol 18:85–100.

Newton, A.H., A. Cardani, and T.J. Braciale. 2016. The host immune response in respiratory virus infection: balancing virus clearance and immunopathology. Semin Immunopathol 38:471–482.

Noval Rivas, M., and M. Arditi. 2023. Kawasaki Disease and Multisystem Inflammatory Syndrome in Children: Common Inflammatory Pathways of Two Distinct Diseases. Rheum Dis Clin North Am 49:647–659.

Patel, J.M. 2022. Multisystem Inflammatory Syndrome in Children (MIS-C). Curr Allergy Asthma Rep 22:53–60.

Pawlowski, K.D., J.T. Duffy, A. Tiwari, M. Zannikou, and I.V. Balyasnikova. 2023. Bi-Specific Killer Cell Engager Enhances NK Cell Activity against Interleukin-13 Receptor Alpha-2 Positive Gliomas. Cells 12:

Qiu, W.Q., D. de Bruin, B.H. Brownstein, R. Pearse, and J.V. Ravetch. 1990. Organization of the human and mouse low-affinity Fc gamma R genes: duplication and recombination. Science 248:732–735.

Rakova, J., I. Truxova, P. Holicek, C. Salek, M. Hensler, L. Kasikova, J. Pasulka, M. Holubova, M. Kovar, D. Lysak, J.P. Kline, Z. Racil, L. Galluzzi, R. Spisek, and J. Fucikova. 2021. TIM-3 levels correlate with enhanced NK cell cytotoxicity and improved clinical outcome in AML patients. Oncoimmunology 10:1889822.

Ramaswamy, A., N.N. Brodsky, T.S. Sumida, M. Comi, H. Asashima, K.B. Hoehn, N. Li, Y. Liu, A. Shah, N.G. Ravindra, J. Bishai, A. Khan, W. Lau, B. Sellers, N. Bansal, P. Guerrerio, A. Unterman, V. Habet, A.J. Rice, J. Catanzaro, H. Chandnani, M. Lopez, N. Kaminski, C.S. Dela Cruz, J.S. Tsang, Z. Wang, X. Yan, S.H. Kleinstein, D. van Dijk, R.W. Pierce, D.A. Hafler, and C.L. Lucas. 2021. Immune dysregulation and autoreactivity correlate with disease severity in SARS-CoV-2-associated multisystem inflammatory syndrome in children. Immunity 54:1083–1095.e1087.

Ravichandran, S., J. Tang, G. Grubbs, Y. Lee, S. Pourhashemi, L. Hussaini, S.A. Lapp, R.C. Jerris, V. Singh, A. Chahroudi, E.J. Anderson, C.A. Rostad, and S. Khurana. 2021. SARS-CoV-2 immune repertoire in MIS-C and pediatric COVID-19. Nat Immunol 22:1452–1464.

Richardson, S.I., N.P. Manamela, B.M. Motsoeneng, H. Kaldine, F. Ayres, Z. Makhado, M. Mennen, S. Skelem, N. Williams, N.J. Sullivan, J. Misasi, G.G. Gray, L.G. Bekker, V. Ueckermann, T.M. Rossouw, M.T. Boswell, N.A.B. Ntusi, W.A. Burgers, and P.L. Moore. 2022. SARS-CoV-2 Beta and Delta variants trigger Fc effector function with increased cross-reactivity. Cell Rep Med 3:100510.

Rodriguez, P.C., A.C. Ochoa, and A.A. Al-Khami. 2017. Arginine Metabolism in Myeloid Cells Shapes Innate and Adaptive Immunity. Front Immunol 8:93.

Rogers, T.F., F. Zhao, D. Huang, N. Beutler, A. Burns, W.-t. He, O. Limbo, C. Smith, G. Song, J. Woehl, L. Yang, R.K. Abbott, S. Callaghan, E. Garcia, J. Hurtado, M. Parren, L. Peng, S. Ramirez, J. Ricketts, M.J. Ricciardi, S.A. Rawlings, N.C. Wu, M. Yuan, D.M. Smith, D. Nemazee, J.R. Teijaro, J.E. Voss, I.A. Wilson, R. Andrabi, B. Briney, E. Landais, D. Sok, J.G. Jardine, and D.R. Burton. 2020. Isolation of potent SARS-CoV-2 neutralizing antibodies and protection from disease in a small animal model. Science 369:956–963.

Rostad, C.A., X. Chen, H.Y. Sun, L. Hussaini, A. Lu, M.A. Perez, H.M. Hsiao, L.J. Anderson, and E.J. Anderson. 2022. Functional antibody responses to SARS-CoV-2 variants in children with COVID-19, MIS-C, and after two doses of BNT162b2 vaccination. J Infect Dis

Rowley, A.H. 2020. Multisystem Inflammatory Syndrome in Children and Kawasaki Disease: Two Different Illnesses with Overlapping Clinical Features. J Pediatr 224:129–132.

Rybkina, K., J.N. Bell, M.C. Bradley, T. Wohlbold, M. Scafuro, W. Meng, R.C. Korenberg, J. Davis-Porada, B.R. Anderson, R.J. Weller, J.D. Milner, A. Moscona, M. Porotto, E.T. Luning Prak, K. Pethe, T.J. Connors, and D.L. Farber. 2023. SARS-CoV-2 infection and recovery in children: Distinct T cell responses in MIS-C compared to COVID-19. J Exp Med 220:

Sacco, K., R. Castagnoli, S. Vakkilainen, C. Liu, O.M. Delmonte, C. Oguz, I.M. Kaplan, S. Alehashemi, P.D. Burbelo, F. Bhuyan, A.A. de Jesus, K. Dobbs, L.B. Rosen, A. Cheng, E. Shaw, M.S. Vakkilainen, F. Pala, J. Lack, Y. Zhang, D.L. Fink, V. Oikonomou, A.L. Snow, C.L. Dalgard, J. Chen, B.A. Sellers, G.A. Montealegre Sanchez, K. Barron, E. Rey-Jurado, C. Vial, M.C. Poli, A. Licari, D. Montagna, G.L. Marseglia, F. Licciardi, U. Ramenghi, V. Discepolo, A. Lo Vecchio, A. Guarino, E.M. Eisenstein, L. Imberti, A. Sottini, A. Biondi, S. Mató, D. Gerstbacher, M. Truong, M.A. Stack, M. Magliocco, M. Bosticardo, T. Kawai, J.J. Danielson, T. Hulett, M. Askenazi, S. Hu, J.I. Cohen, H.C. Su, D.B. Kuhns, M.S. Lionakis, T.M. Snyder, S.M. Holland, R. Goldbach-Mansky, J.S. Tsang, and L.D. Notarangelo. 2022. Immunopathological signatures in multisystem inflammatory syndrome in children and pediatric COVID-19. Nat Med 28:1050–1062.

Sayers, E.W., E.E. Bolton, J.R. Brister, K. Canese, J. Chan, D.C. Comeau, R. Connor, K. Funk, C. Kelly, S. Kim, T. Madej, A. Marchler-Bauer, C. Lanczycki, S. Lathrop, Z. Lu, F. Thibaud-Nissen, T. Murphy, L. Phan, Y. Skripchenko, T. Tse, J. Wang, R. Williams, B.W. Trawick, K.D. Pruitt, and S.T. Sherry. 2022. Database resources of the national center for biotechnology information. Nucleic Acids Res 50:D20–d26.

Schäfer, A., F. Muecksch, J.C.C. Lorenzi, S.R. Leist, M. Cipolla, S. Bournazos, F. Schmidt, R.M. Maison, A. Gazumyan, D.R. Martinez, R.S. Baric, D.F. Robbiani, T. Hatziioannou, J.V. Ravetch, P.D. Bieniasz, R.A. Bowen, M.C. Nussenzweig, and T.P. Sheahan. 2021. Antibody potency, effector function, and combinations in protection and therapy for SARS-CoV-2 infection in vivo. J Exp Med 218:

Schoeman, D., and B.C. Fielding. 2019. Coronavirus envelope protein: current knowledge. Virology Journal 16:69.

Sharma, C., M. Ganigara, C. Galeotti, J. Burns, F.M. Berganza, D.A. Hayes, D. Singh-Grewal, S. Bharath, S. Sajjan, and J. Bayry. 2021. Multisystem inflammatory syndrome in children and Kawasaki disease: a critical comparison. Nat Rev Rheumatol 17:731–748.

Shim, S., S. Lee, Y. Hisham, S. Kim, T.T. Nguyen, A.S. Taitt, J. Hwang, H. Jhun, H.Y. Park, Y. Lee, S.C. Yeom, S.Y. Kim, Y.G. Kim, and S. Kim. 2022. Comparison of the Seven Interleukin-32 Isoforms’ Biological Activities: IL-32θ Possesses the Most Dominant Biological Activity. Front Immunol 13:837588.

Shu, Y., and J. McCauley. 2017. GISAID: Global initiative on sharing all influenza data - from vision to reality. Euro Surveill 22:

Sönmez, H.E., Ş. Çağlayan, G. Otar Yener, E.Z. Başar, K. Ulu, M. Çakan, V. Guliyeva, E. Bağlan, K. Öztürk, D. Demirkol, F. Demir, G. Karadağ Ş, S. Özdel, N. Aktay Ayaz, and B. Sözeri. 2022. The Multifaceted Presentation of the Multisystem Inflammatory Syndrome in Children: Data from a Cluster Analysis. J Clin Med 11:

Stranneheim, H., K. Lagerstedt-Robinson, M. Magnusson, M. Kvarnung, D. Nilsson, N. Lesko, M. Engvall, B.-M. Anderlid, H. Arnell, C.B. Johansson, M. Barbaro, E. Björck, H. Bruhn, J. Eisfeldt, C. Freyer, G. Grigelioniene, P. Gustavsson, A. Hammarsjö, M. Hellström-Pigg, E. Iwarsson, A. Jemt, M. Laaksonen, S.L. Enoksson, H. Malmgren, K. Naess, M. Nordenskjöld, M. Oscarson, M. Pettersson, C. Rasi, A. Rosenbaum, E. Sahlin, E. Sardh, T. Stödberg, B. Tesi, E. Tham, H. Thonberg, V. Töhönen, U. von Döbeln, D. Vassiliou, S. Vonlanthen, A.-C. Wikström, J. Wincent, O. Winqvist, A. Wredenberg, S. Ygberg, R.H. Zetterström, P. Marits, M.J. Soller, A. Nordgren, V. Wirta, A. Lindstrand, and A. Wedell. 2021. Integration of whole genome sequencing into a healthcare setting: high diagnostic rates across multiple clinical entities in 3219 rare disease patients. Genome Medicine 13:40.

Szabo, P.A., P. Dogra, J.I. Gray, S.B. Wells, T.J. Connors, S.P. Weisberg, I. Krupska, R. Matsumoto, M.M.L. Poon, E. Idzikowski, S.E. Morris, C. Pasin, A.J. Yates, A. Ku, M. Chait, J. Davis-Porada, X.V. Guo, J. Zhou, M. Steinle, S. Mackay, A. Saqi, M.R. Baldwin, P.A. Sims, and D.L. Farber. 2021. Longitudinal profiling of respiratory and systemic immune responses reveals myeloid cell-driven lung inflammation in severe COVID-19. Immunity 54:797–814.e796.

Tay, M.Z., K. Wiehe, and J. Pollara. 2019. Antibody-Dependent Cellular Phagocytosis in Antiviral Immune Responses. Frontiers in Immunology 10:

Thiriard, A., B. Meyer, C.S. Eberhardt, N. Loevy, S. Grazioli, W. Adouan, P. Fontannaz, F. Marechal, A.G. L’Huillier, C.A. Siegrist, D. Georges, A. Putignano, A. Marchant, A.M. Didierlaurent, and G. Blanchard-Rohner. 2023. Antibody response in children with multisystem inflammatory syndrome related to COVID-19 (MIS-C) compared to children with uncomplicated COVID-19. Front Immunol 14:1107156.

Thomas, S. 2020. The Structure of the Membrane Protein of SARS-CoV-2 Resembles the Sugar Transporter SemiSWEET. Pathog Immun 5:342–363.

Troyano-Hernáez, P., R. Reinosa, and Á. Holguín. 2021. Evolution of SARS-CoV-2 Envelope, Membrane, Nucleocapsid, and Spike Structural Proteins from the Beginning of the Pandemic to September 2020: A Global and Regional Approach by Epidemiological Week. Viruses 13:

Ullah, I., J. Prévost, M.S. Ladinsky, H. Stone, M. Lu, S.P. Anand, G. Beaudoin-Bussières, K. Symmes, M. Benlarbi, S. Ding, R. Gasser, C. Fink, Y. Chen, A. Tauzin, G. Goyette, C. Bourassa, H. Medjahed, M. Mack, K. Chung, C.B. Wilen, G.A. Dekaban, J.D. Dikeakos, E.A. Bruce, D.E. Kaufmann, L. Stamatatos, A.T. McGuire, J. Richard, M. Pazgier, P.J. Bjorkman, W. Mothes, A. Finzi, P. Kumar, and P.D. Uchil. 2021a. Live Imaging of SARS-CoV-2 Infection in Mice Reveals Neutralizing Antibodies Require Fc Function for Optimal Efficacy. bioRxiv

Ullah, I., J. Prévost, M.S. Ladinsky, H. Stone, M. Lu, S.P. Anand, G. Beaudoin-Bussières, K. Symmes, M. Benlarbi, S. Ding, R. Gasser, C. Fink, Y. Chen, A. Tauzin, G. Goyette, C. Bourassa, H. Medjahed, M. Mack, K. Chung, C.B. Wilen, G.A. Dekaban, J.D. Dikeakos, E.A. Bruce, D.E. Kaufmann, L. Stamatatos, A.T. McGuire, J. Richard, M. Pazgier, P.J. Bjorkman, W. Mothes, A. Finzi, P. Kumar, and P.D. Uchil. 2021b. Live imaging of SARS-CoV-2 infection in mice reveals that neutralizing antibodies require Fc function for optimal efficacy. Immunity 54:2143–2158.e2115.

Vahidy, F.S., J.C. Nicolas, J.R. Meeks, O. Khan, A. Pan, S.L. Jones, F. Masud, H.D. Sostman, R. Phillips, J.D. Andrieni, B.A. Kash, and K. Nasir. 2020. Racial and ethnic disparities in SARS-CoV-2 pandemic: analysis of a COVID-19 observational registry for a diverse US metropolitan population. BMJ Open 10:e039849.

Vallera, D.A., S. Ferrone, B. Kodal, P. Hinderlie, L. Bendzick, B. Ettestad, C. Hallstrom, N.A. Zorko, A. Rao, N. Fujioka, C.J. Ryan, M.A. Geller, J.S. Miller, and M. Felices. 2020. NK-Cell-Mediated Targeting of Various Solid Tumors Using a B7-H3 Tri-Specific Killer Engager In Vitro and In Vivo. Cancers 12:2659.

Vallera, D.a.M., JS. 2017. THERAPEUTIC COMPOUNDS AND METHODS. In.

Villanueva, J., S. Lee, E.H. Giannini, T.B. Graham, M.H. Passo, A. Filipovich, and A.A. Grom. 2005. Natural killer cell dysfunction is a distinguishing feature of systemic onset juvenile rheumatoid arthritis and macrophage activation syndrome. Arthritis Res Ther 7:R30–37.

Vogel, T.P., K.A. Top, C. Karatzios, D.C. Hilmers, L.I. Tapia, P. Moceri, L. Giovannini-Chami, N. Wood, R.E. Chandler, N.P. Klein, E.P. Schlaudecker, M.C. Poli, E. Muscal, and F.M. Munoz. 2021. Multisystem inflammatory syndrome in children and adults (MIS-C/A): Case definition & guidelines for data collection, analysis, and presentation of immunization safety data. Vaccine 39:3037–3049.

Waterhouse, A.M., J.B. Procter, D.M. Martin, M. Clamp, and G.J. Barton. 2009. Jalview Version 2--a multiple sequence alignment editor and analysis workbench. Bioinformatics 25:1189–1191.

Way, J., Burrill, DR, Hof, KS. 2017. HUMANIZED GLYCOPHORIN A ANTIBODIES AND USES THEREOF. In W.I.P. Organization, editor United States. 110.

Weisberg, S.P., T.J. Connors, Y. Zhu, M.R. Baldwin, W.H. Lin, S. Wontakal, P.A. Szabo, S.B. Wells, P. Dogra, J. Gray, E. Idzikowski, D. Stelitano, F.T. Bovier, J. Davis-Porada, R. Matsumoto, M.M.L. Poon, M. Chait, C. Mathieu, B. Horvat, D. Decimo, K.E. Hudson, F.D. Zotti, Z.C. Bitan, F. La Carpia, S.A. Ferrara, E. Mace, J. Milner, A. Moscona, E. Hod, M. Porotto, and D.L. Farber. 2021. Distinct antibody responses to SARS-CoV-2 in children and adults across the COVID-19 clinical spectrum. Nat Immunol 22:25–31.

Whittaker, E., A. Bamford, J. Kenny, M. Kaforou, C.E. Jones, P. Shah, P. Ramnarayan, A. Fraisse, O. Miller, P. Davies, F. Kucera, J. Brierley, M. McDougall, M. Carter, A. Tremoulet, C. Shimizu, J. Herberg, J.C. Burns, H. Lyall, and M. Levin. 2020. Clinical Characteristics of 58 Children With a Pediatric Inflammatory Multisystem Syndrome Temporally Associated With SARS-CoV-2. Jama 324:259–269.

Winkler, E.S., P. Gilchuk, J. Yu, A.L. Bailey, R.E. Chen, Z. Chong, S.J. Zost, H. Jang, Y. Huang, J.D. Allen, J.B. Case, R.E. Sutton, R.H. Carnahan, T.L. Darling, A.C.M. Boon, M. Mack, R.D. Head, T.M. Ross, J.E. Crowe, Jr., and M.S. Diamond. 2021. Human neutralizing antibodies against SARS-CoV-2 require intact Fc effector functions for optimal therapeutic protection. Cell 184:1804–1820.e1816.

Wu, J., F.X. Gao, C. Wang, M. Qin, F. Han, T. Xu, Z. Hu, Y. Long, X.M. He, X. Deng, D.L. Ren, and T.Y. Dai. 2019. IL-6 and IL-8 secreted by tumour cells impair the function of NK cells via the STAT3 pathway in oesophageal squamous cell carcinoma. J Exp Clin Cancer Res 38:321.

Wu, J., Y. Li, A. Rendahl, and M. Bhargava. 2022. Novel Human FCGR1A Variants Affect CD64 Functions and Are Risk Factors for Sarcoidosis. Front Immunol 13:841099.

Wu, J., R. Lin, J. Huang, W. Guan, W.S. Oetting, P. Sriramarao, and M.N. Blumenthal. 2014. Functional Fcgamma receptor polymorphisms are associated with human allergy. PLoS One 9:e89196.

Yamin, R., A.T. Jones, H.H. Hoffmann, A. Schäfer, K.S. Kao, R.L. Francis, T.P. Sheahan, R.S. Baric, C.M. Rice, J.V. Ravetch, and S. Bournazos. 2021. Fc-engineered antibody therapeutics with improved anti-SARS-CoV-2 efficacy. Nature 599:465–470.

Yasuhara, J., K. Watanabe, H. Takagi, N. Sumitomo, and T. Kuno. 2021. COVID-19 and multisystem inflammatory syndrome in children: A systematic review and meta-analysis. Pediatr Pulmonol 56:837–848.

Yasui, F., M. Kohara, M. Kitabatake, T. Nishiwaki, H. Fujii, C. Tateno, M. Yoneda, K. Morita, K. Matsushima, S. Koyasu, and C. Kai. 2014. Phagocytic cells contribute to the antibody-mediated elimination of pulmonary-infected SARS coronavirus. Virology 454-455:157–168.

Zhang, A., H.D. Stacey, M.R. D’Agostino, Y. Tugg, A. Marzok, and M.S. Miller. 2023. Beyond neutralization: Fc-dependent antibody effector functions in SARS-CoV-2 infection. Nat Rev Immunol 23:381–396.

Zhang, A., H.D. Stacey, M.R. D’Agostino, Y. Tugg, A. Marzok, and M.S. Miller. 2022. Beyond neutralization: Fc-dependent antibody effector functions in SARS-CoV-2 infection. Nature Reviews Immunology

