## Supplemental Figures for "Antibody-mediated cellular responses are dysregulated in Multisystem Inflammatory Syndrome in Children (MIS-C)"

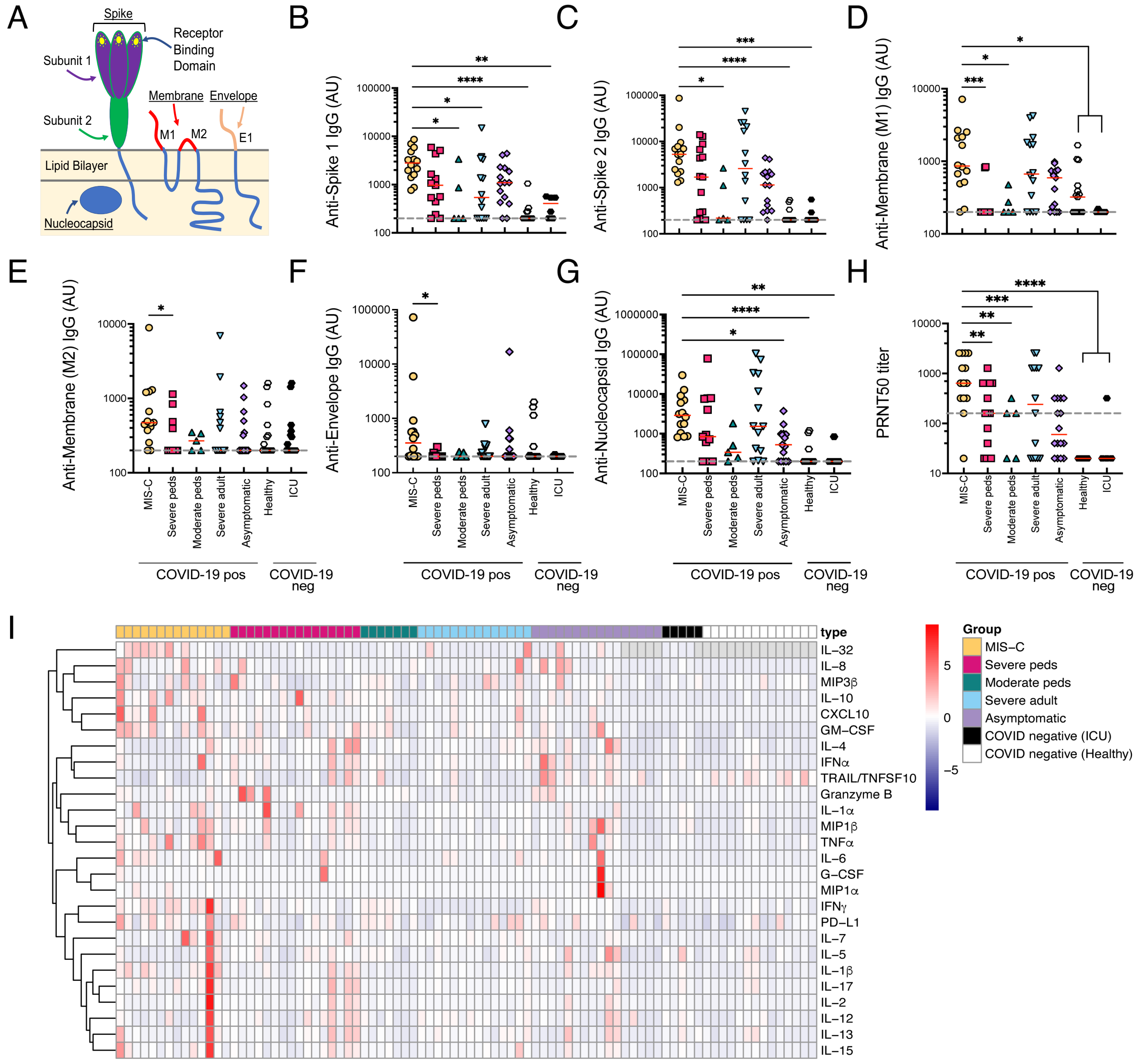


**Supplemental Figure 1. Profiling of SARS-CoV-2 specific and neutralization antibody and cytokine responses. (A)** Schematic of SARS-CoV-2 viruses indicating which targets on the virus that the antibody responses were against. (**B-G**) SARS-CoV-2 antibody reactivities stratified by group; MIS-C (n=14), severe peds (n = 13), moderate peds (n=5), severe adult (n = 14), asymptomatic (n=15), COVID-19 negative healthy (n=15), COVID-19 negative ICU (n=6).

Patients that received convalescent plasma were removed from this analysis. Graphs depict IgG titers against SARS-CoV-2 **(B)** Spike subunit 1 (S1), **(C)** Spike subunit 2 (S2), **(D-E)** membrane protein (M1 and M2), and **(F)** envelope protein peptides that are expressed on the surface of SARS-CoV-2 and **(G)** nucleocapsid protein which is internal for infectious viruses. Red lines depict the median. Dashed line is the limit of detection. (**H**) 50% plaque reduction neutralization test (PRNT50) to SARS-CoV-2 isolate 2019-nCoV/USA-WA1/2020, stratified by group. Statistical analyses were performed using a Kruskal-Wallis test with Dunn's multiple comparison test. Red lines depict median. *, P <0.05, ** P< 0.01. ***P<0.001, **** P < 0.0001. **(I)** Profiles of cytokines and chemokines in plasma shown as a heatmap, stratified by group and divided out by individual subject; MIS-C (n=14), severe pediatrics (peds) (n = 16), moderate pediatrics (n=7), severe adult (n = 14), asymptomatic pediatrics (n=15), COVID-19 negative healthy pediatrics (n=15), COVID-19 negative ICU pediatrics (n=6). Color intensity of each rectangle represents the average Z-score of cytokine measures within each group.

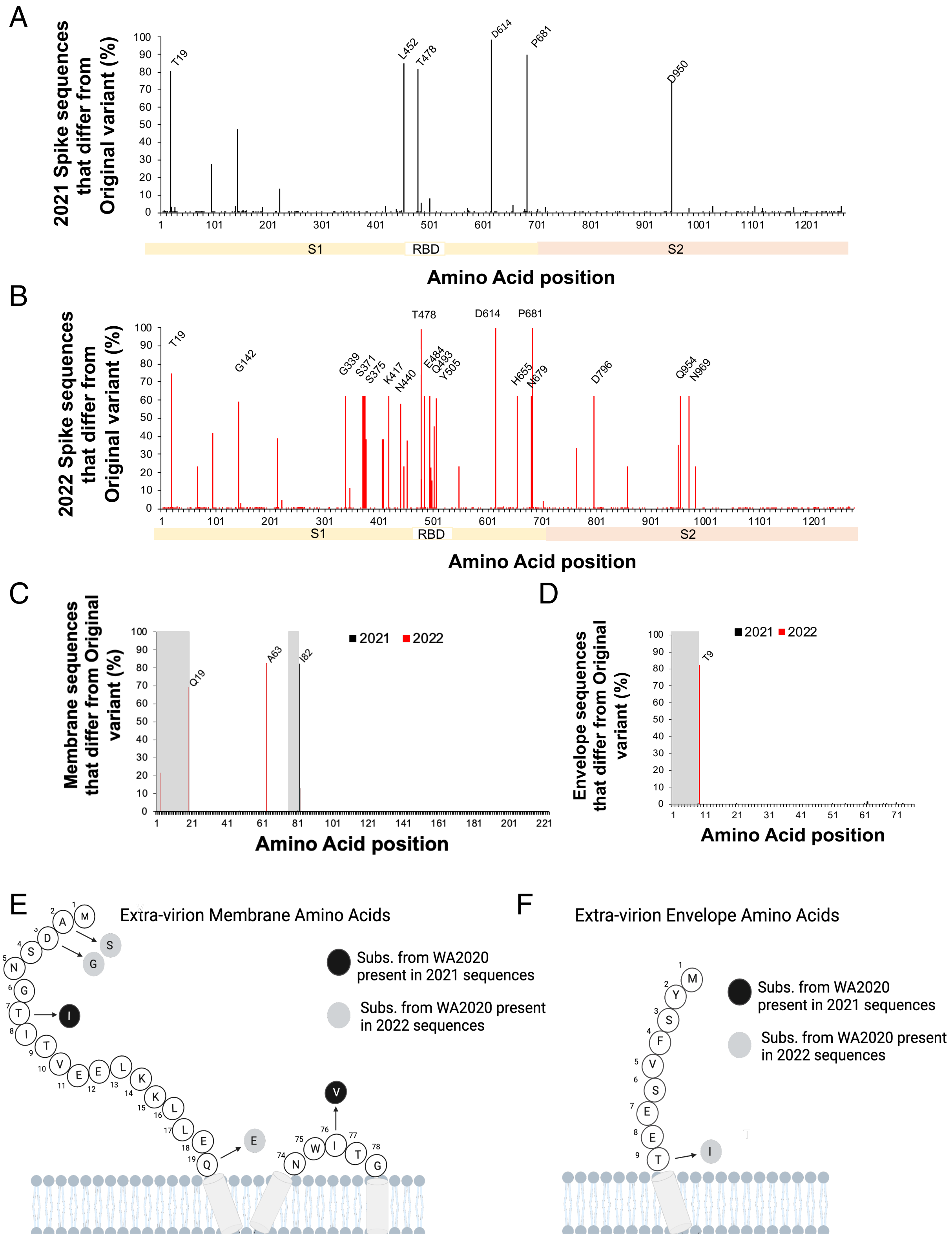
A
2021 Spike sequences that differ from Original variant (%)
T19
L452
T478
D614
P681
D950
S1
RBD
S2
Amino Acid position
B
2022 Spike sequences that differ from Original variant (%)
T19
G142
G339
S371
S375
K417
N440
T478
E484
Q493
Y505
D614
H655
N679
P681
D796
Q954
N969
S1
RBD
S2
Amino Acid position
C
Membrane sequences that differ from Original variant (%)
2021
2022
Q19
A63
I82
Amino Acid position
D
Envelope sequences that differ from Original variant (%)
2021
2022
T9
Amino Acid position
E
Extra-virion Membrane Amino Acids
Subs. from WA2020 present in 2021 sequences
Subs. from WA2020 present in 2022 sequences
F
Extra-virion Envelope Amino Acids
Subs. from WA2020 present in 2021 sequences
Subs. from WA2020 present in 2022 sequences

**Supplemental Figure 2. Sequence variation analysis of protein and peptides on the surface of SARS-CoV-2. (A-B)** Percentage of analyzed SARS-CoV-2 spike gene sequences differing from the original SARS-CoV-2 strain by amino acid position in 2021 **(A)** and 2022 **(B).** Functionally significant domains (S1, S2 and receptor binding motif or RBD) are indicated by shaded bars**. (C-D)** Percentage of analyzed SARS-CoV-2 membrane **(C)** and envelope **(D)** sequences differing from the original SARS-CoV-2 strain by amino acid position in 2021 and 2022. Sequences collected in 2021 are depicted in black and those collected in 2022 are depicted in red. Amino acid substitutions present at a frequency greater than 50% in their given year are labeled on the graph. The predicted extracellular domains are indicated by gray shading. **(E-F)** Schematic diagram of the SARS-CoV-2 membrane **(E)** and envelope **(F)** transmembrane proteins illustrating the predicted topology and sites of sequence variability present in genome sequences in the GISAID hCoV-19 Variant database relative to the original reference strain. The individual amino acid residues for predicted extracellular portions of the original strain are depicted with open circles. Amino acid substitutions present at ≥0.2% frequency among 2021 variants are depicted as black circles and those among 2022 variants are depicted as grey circles.

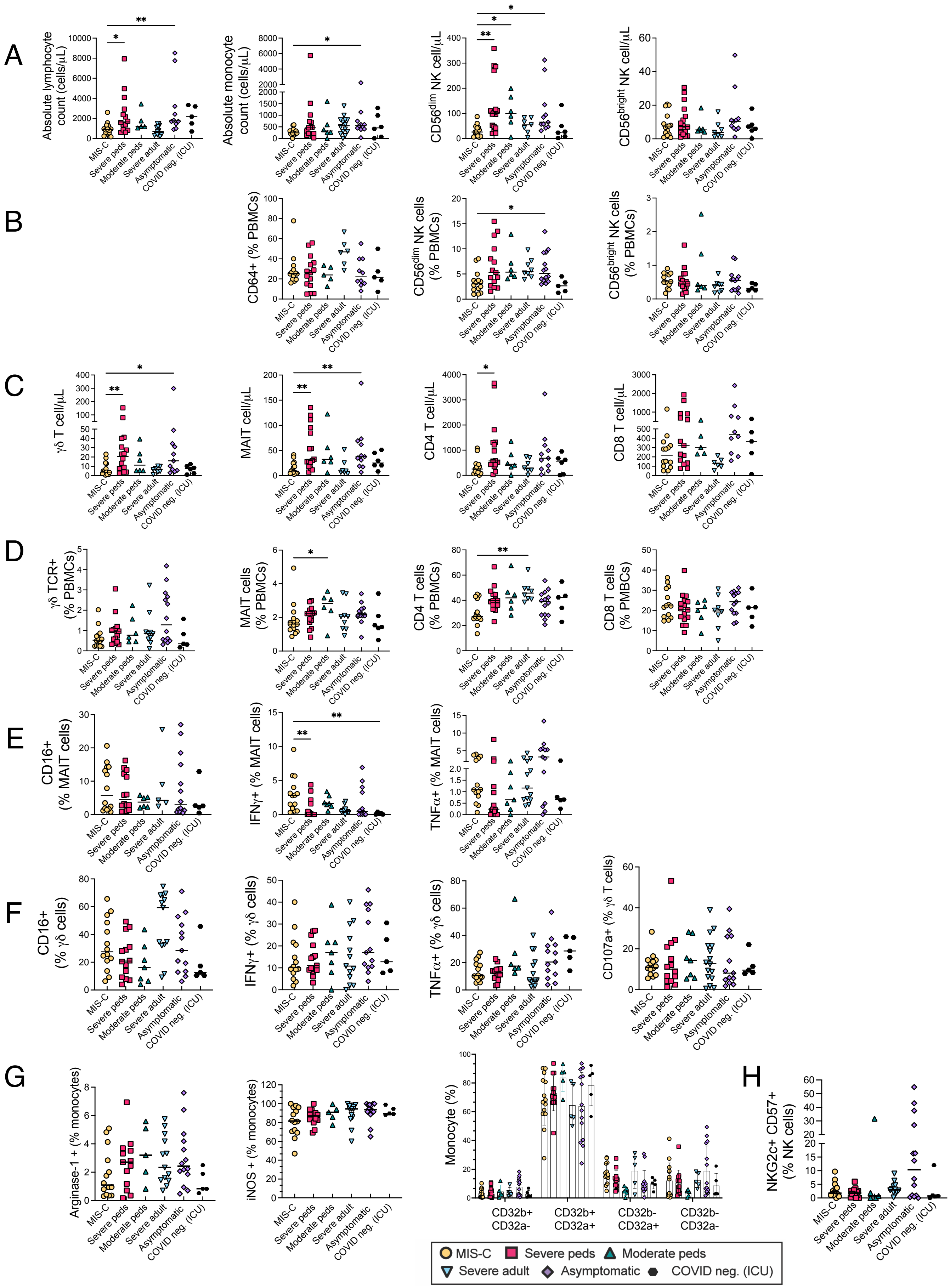
A
Absolute lymphocyte count (cells/μL)
Absolute monocyte count (cells/μL)
CD56dim NK cell/μL
CD56bright NK cell/μL
B
CD64+ (% PBMCs)
CD56dim NK cells (% PBMCs)
CD56bright NK cells (% PBMCs)
C
γδ T cell/μL
MAIT cell/μL
CD4 T cell/μL
CD8 T cell/μL
D
γδ TCR+ (% PBMCs)
MAIT cells (% PBMCs)
CD4 T cells (% PBMCs)
CD8 T cells (% PMBCs)
E
CD16+ (% MAIT cells)
IFNγ+ (% MAIT cells)
TNFα+ (% MAIT cells)
F
CD16+ (% γδ cells)
IFNγ+ (% γδ cells)
TNFα+ (% γδ cells)
CD107a+ (% γδ T cells)
G
Arginase-1 + (% monocytes)
iNOS + (% monocytes)
Monocyte (%)
CD32b+ CD32a-
CD32b+ CD32a+
CD32b- CD32a+
CD32b- CD32a-
H
NKG2c+ CD57+ (% NK cells)
MIS-C
Severe peds
Moderate peds
Severe adult
Asymptomatic
COVID neg. (ICU)

**Supplemental Figure 3 Cell counts differ between MIS-C and acute COVID-19 patients, but monocytes, γδ T cells and MAIT cells are phenotypically similar between groups. (A-B)** Cell counts **(A)** and precent **(B)** of monocytes, lymphocytes, CD56dim and CD56bright NK cells in the blood. **(C-D).** Cell counts **(C)** in the blood and percent of PBMCs **(D)** of γδ T cells, MAIT cells, CD4 T cells and CD8 T cells. **(E-F)** Phenotypic and functional markers including CD16, IFNγ, TNFα, and degranulation on MAIT cells **(E)** and γδ T cells **(F)** in an ADCC assay**. (G)** Markers of monocyte subsets (iNOS, Arginase- 1, CD32a and CD32b). **(H)** NKG2c+/CD57+ NK cells. All data is stratified by group. Statistical analyses were performed using a Kruskal-Wallis test with Dunn's multiple comparison test. Black lines depict median. *, $P < 0.05$, ** $P < 0.01$. ***$P < 0.001$, **** $P < 0.0001$.

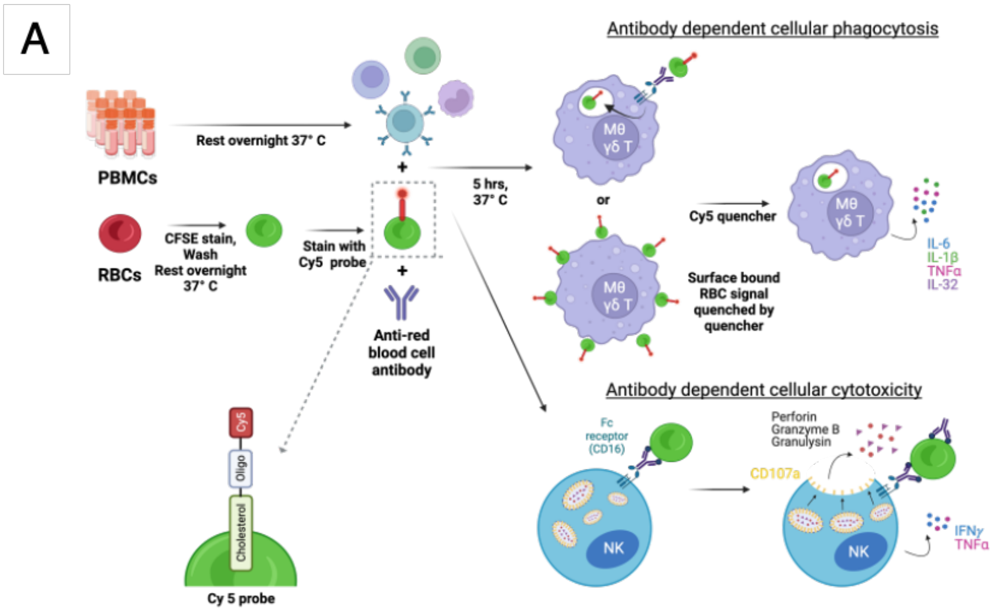
A
Antibody dependent cellular phagocytosis
PBMCs
Rest overnight 37° C
RBCs
CFSE stain, Wash Rest overnight 37° C
Stain with Cy5 probe
Anti-red blood cell antibody
5 hrs, 37° C
or
Cy5 quencher
Surface bound RBC signal quenched by quencher
IL-6
IL-1β
TNFα
IL-32
Antibody dependent cellular cytotoxicity
Fc receptor (CD16)
Perforin
Granzyme B
Granulysin
CD107a
NK
IFNγ
TNFα
Cy5
Oligo
Cholesterol
Cy 5 probe

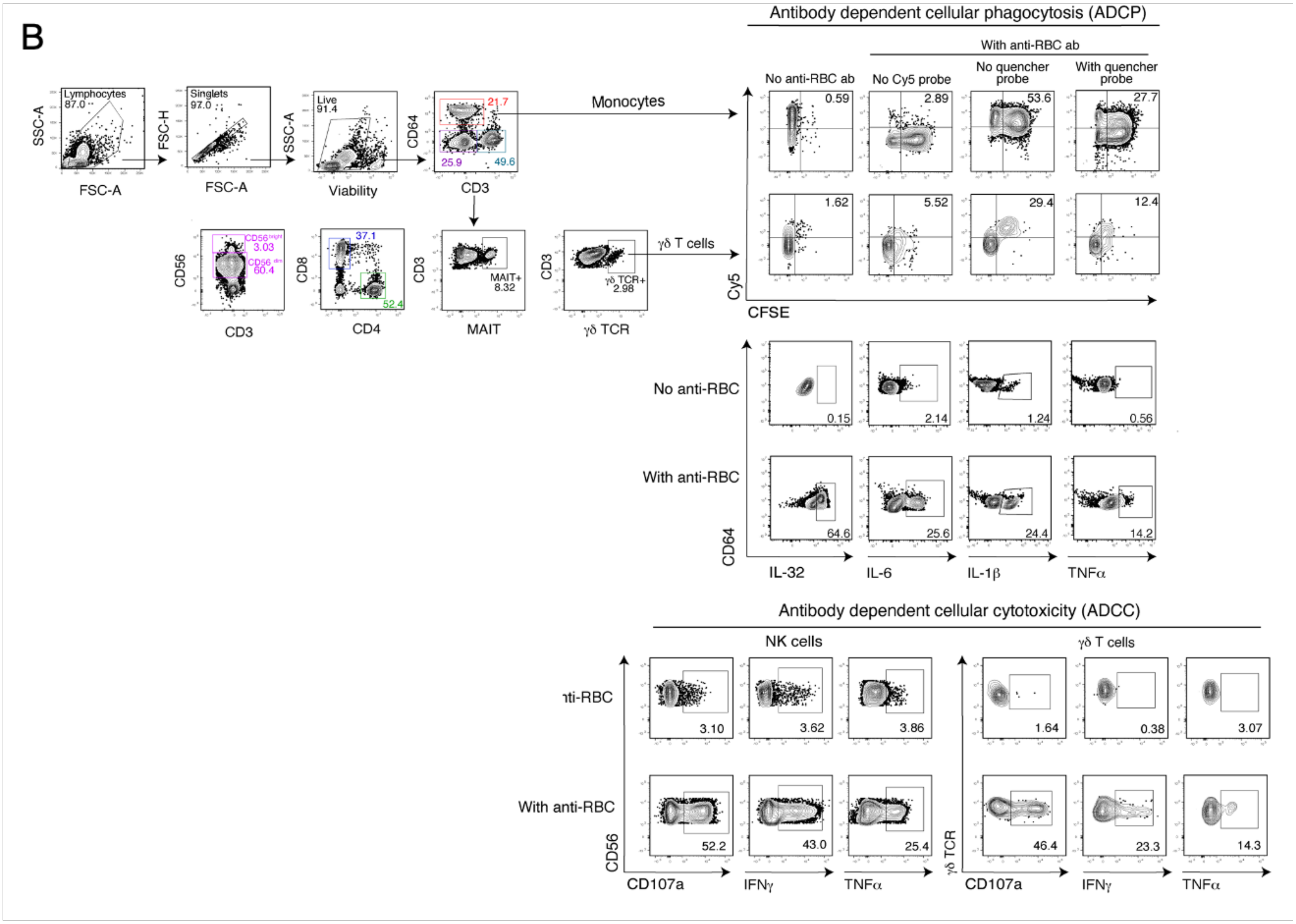
B
Antibody dependent cellular phagocytosis (ADCP)
With anti-RBC ab
No anti-RBC ab
No Cy5 probe
No quencher probe
With quencher probe
Monocytes
γδ T cells
Lymphocytes
Singlets
Live
Cy5
CFSE
No anti-RBC
With anti-RBC
CD64
IL-32
IL-6
IL-1β
TNFα
Antibody dependent cellular cytotoxicity (ADCC)
NK cells
γδ T cells
With anti-RBC
CD56
CD107a
IFNγ
TNFα
γδ TCR

**Supplemental Figure 4. Gating strategies for ADCC and ADCP flow cytometry assay (A)** Schematic of antibody dependent cellular phagocytosis (ADCP) and antibody dependent cellular cytotoxicity (ADCC) assays. For background on the Cy5-oligo-cholesterol probe, the cholesterol part's purpose is to intercalate into the cell membrane to bind the probe to the RBC. The Cy5 allows for the detection of the probe via flow cytometry in the APC channel. The oligo allows for the quenching of the probe by adding a separate probe that is the reverse complement of the original oligo except now has a quencher chemical on one end (quencher reverse complement oligo) that quenches the Cy5 signal (timing of this is later in the protocol). When this quencher reverse complement oligo is added, any phagocyte surface bound RBCs that are not phagocytosed will have their signal quenched, but the RBCs that were phagocytosed completely will not be able to be quenched. *Therefore, the monocytes that have fully phagocytosed RBCs will be counted in the flow cytometry assay as CFSE+ and Cy5+* (Fig. S1).

**(B)** Gating strategy used for phenotyping and functional assays of innate immune cell from concatenated flow cytometry files. Numbers on flow cytometry plots represent % of cells within the gate. FSC, forward scatter. SSC, side scatter.

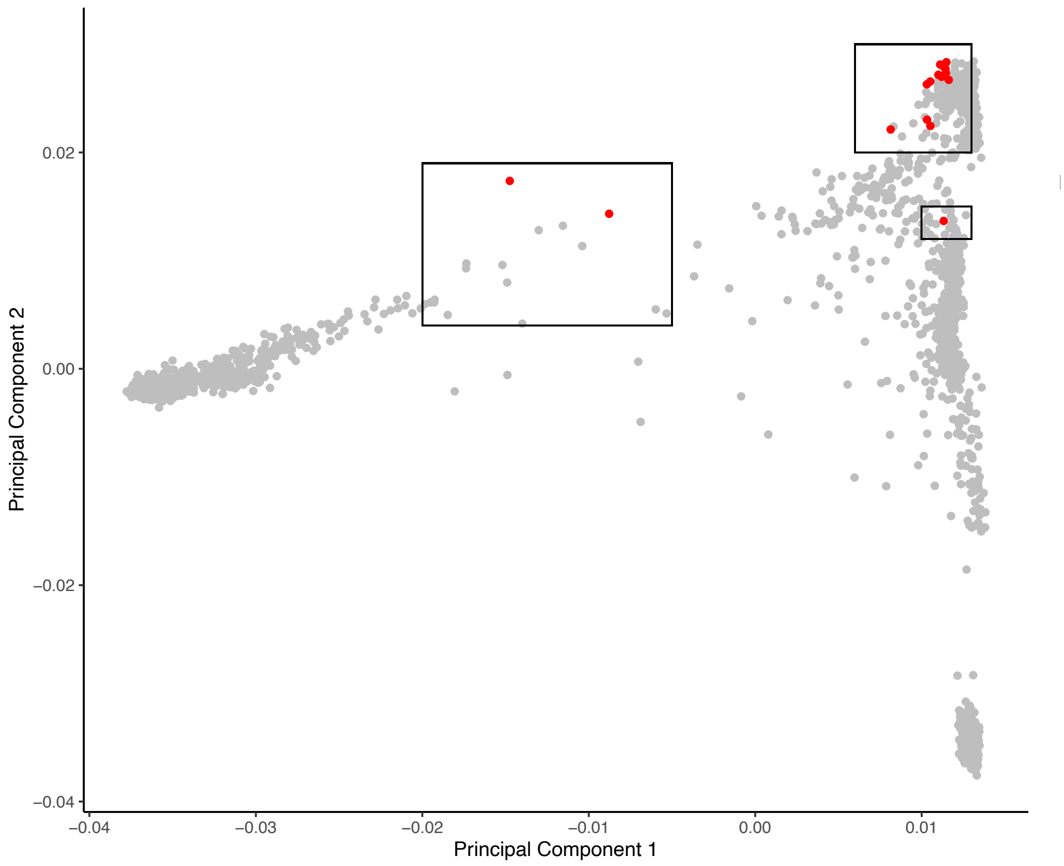


**Supplemental Figure 5. Principal component analysis to compare ethnicities of individuals with MIS-C compared to healthy controls from the 1000 Genomes Project**. Principal component analysis (PCA) was performed on patient genomes and the first two mapped against the 1000 Genomes Project (1kGP). 1kGP participants with proximal ancestry PCs were selected for further genotype comparisons, marked by boxes. The red dots indicate children with MIS-C.

**Supplemental Table 2.** Allele frequencies in Fc receptors between individuals with MIS-C and healthy controls.

| **Fc receptor** | **Variant ID** | **MIS-C (n =20)** | **Healthy controls (n=184)** | **Enrichment in patients** |
|---|---|---|---|---|
| **FCGRIA** | rs1848781 (c.-131 C>G)<br>Prevalence of G allele<br><br>Expression level: G>C | 5% | 0.27% | 18.4 |
| **FCGRIA** | rs587598788 (c.845-23-delTCTTTG)<br>Prevalence of Del<br><br>Expression level: In>Del | 2.5% | 1.6% | 1.53 |
| **FCGRIA** | rs1050204 (c.970 G>A)<br>Prevalence of A allele<br><br>IgG binding capacity: A>G | 0% | 0% | N/A |
| **FCGRIIA** | rs1801274 (c.131 G>A)<br>Prevalence of A allele<br><br>IgG binding capacity: A>G | 47.5% | 52.9% | 0.89 |
| **FCGRIIB** | rs1050501 (c.775 T>C)<br>Prevalence of C<br><br>Inhibitory function: T>C | 10% | 13% | 0.751 |
| **FCGRIIIA** | rs10127939 (c.230 T>A)<br>Prevalence of A<br><br>IgG binding capacity A>T | 25% | 33.7% | 0.742 |
| | rs396991 (c.559 T>G)<br>Prevalence of T<br><br>IgG binging capacity: G>T | 10% | 11.4% | 0.876 |
